# Deep spatial proteomics of ovarian cancer precursor lesions delineates early disease changes and cell-of-origin signatures

**DOI:** 10.1101/2025.03.19.643504

**Authors:** Anuar Makhmut, Mihnea P. Dragomir, Sonja Fritzsche, Markus Moebs, Wolfgang D. Schmitt, Eliane T. Taube, Fabian Coscia

**Affiliations:** Max-Delbrück-Center for Molecular Medicine in the Helmholtz Association (MDC), Spatial Proteomics Group, Berlin, Germany; Charité – Universitätsmedizin Berlin, corporate member of Freie Universität Berlin and Humboldt-Universität zu Berlin, Institute of Pathology, Berlin, Germany; Berlin Institute of Health at Charité – Universitätsmedizin Berlin, Berlin, Germany; German Cancer Consortium (DKTK), Partner Site Berlin, German Cancer Research Center (DKFZ), Heidelberg, Germany; Humboldt-Universität zu Berlin, Institute of Biology, Berlin, Germany; Institute of Pathology, Diagnostik Ernst von Bermann, Potsdam, Germany

## Abstract

High-grade serous ovarian cancer (HGSOC) is a devastating disease that is frequently detected at an incurable stage. Advances in ultrasensitive mass spectrometry-based spatial proteomics have provided a unique opportunity to uncover early molecular events in tumorigenesis and common dysregulated pathways with high therapeutic potential. Here, we present a comprehensive proteomic analysis of serous tubal intraepithelial carcinoma (STIC), the HGSOC precursor lesion, covering more than 10,000 proteins. We found that STICs and concurrent invasive carcinomas were indistinguishable at the global proteomic level, revealing a similar level of molecular heterogeneity. Using cell-type resolved tissue proteomics, we revealed strong cell-of-origin signatures preserved in STICs and invasive tumors and identified early dysregulated pathways of therapeutic relevance, such as an onco-metabolic increase in cholesterol biosynthesis. Finally, we uncovered substantial remodeling of the co-evolving tumor microenvironment, affecting approximately one-third of the stromal proteome, and derived a common signature associated with progressive immunosuppression and extracellular matrix restructuring. In summary, our study highlights the power of spatially resolved quantitative proteomics to dissect the molecular underpinnings of early carcinogenesis and provides a rich proteomic resource for future biomarker and drug target research in ovarian cancer.

## Introduction

Among ovarian cancer subtypes, high-grade serous ovarian carcinoma (HGSOC) stands out as the variant that causes the highest number of fatalities related to this malignancy^1^. It is often diagnosed at an advanced stage and is characterized by frequent peritoneal metastasis and tumor recurrence following platinum-based chemotherapy. Genomic analyses have identified key genetic features of HGSOC, including pronounced genomic instability and vast copy number alterations, few recurrent mutations other than ubiquitous *TP53* mutations ^2^, homologous recombination (HRD) deficiency in roughly half of the tumors, often caused by *BRCA1/2* germline and somatic mutations, and high inter- and intra-tumoral heterogeneity ^3^. Moreover, several studies have highlighted the polyclonal nature of HGSOC and its pronounced spatial and temporal heterogeneity ^4–6^. Mass spectrometry-based proteomics studies have addressed how HGSOC genomic alterations translate to the protein level, identifying dysregulated pathways associated with patient survival ^7^ and chemotherapy response ^8,9^. These studies also revealed that the four bulk transcriptomic subtypes (mesenchymal, proliferative, immunoreactive, and differentiated) are reflected in the proteome^7^.

Despite these major efforts to map the proteogenomic disease landscape, far less understood are HGSOC precursor lesions, which are histologically present in 20-60% of all HGSOC patients ^10,11^. Termed serous tubal intraepithelial carcinomas (STICs), these lesions share histological, molecular and genetic features with advanced HGSOC. Although it was first somewhat surprising to find the precursor for ovarian carcinoma in the tubal epithelium, the concept of STICs is widely acknowledged ^12–14^. Although STICs were previously profiled by whole-exome sequencing, which showed that they already present high genome instability and pronounced copy number alterations ^14,15^, very little is known about their proteomic makeup. In particular, we lack a detailed understanding of the extent to which precursor lesions molecularly diverge from normal fallopian tube epithelial cells to ultimately form morphologically distinct invasive tumors. As protein abundance is directly related to cellular phenotype and function ^16^, unraveling precancerous proteome states can provide important insights into the earliest stages of HGSOC development and progression. Such knowledge is of paramount importance for preventive medicine approaches and for identifying protein-based drug targets that are likely to be present in the majority of tumor clones. In addition, we lack a deeper understanding of the co-evolving tumor microenvironment (TME), comprising the extracellular matrix (ECM) and diverse stromal and immune cell types, which are critically involved in all phases of HGSOC development ^17^. Several lines of evidence support the crucial role of the HGSOC TME in regulating disease progression ^18,19^ and immunosuppression ^20^, which dictate therapeutic outcomes ^21^. Our own work recently identified a stromal signature of ovarian cancer metastasis and identified nicotinamide N-methyltransferase as a promising new drug target against cancer-associated fibroblasts ^22^.

Here, we performed deep spatial proteomics of laser-microdissected fallopian tube epithelial cells, STICs, and concurrent invasive carcinomas, as well as their adjacent stromal regions. We present the first comprehensive proteomic map of ovarian cancer precursor lesions, encompassing more than 10,000 proteins in 36 patients. Our data revealed strong cell-of-origin proteome signatures preserved in STICs and concurrent invasive tumors, nominated onco-metabolic adaptations as early events during HGSOC development, and dissected the progressive co-evolution of the HGSOC tumor microenvironment.

## Results

### Histopathology-guided ultra-low input proteomics of HGSOC precursor lesions

To study the cell type- and compartment-resolved proteomic progression of HGSOC precursor lesions (**Fig. 1A**), we selected a cohort of 36 patients with histologically confirmed serous tubal intraepithelial carcinoma (STIC). Precursor lesions were classified according to the criteria proposed by Vang *et al*. ^23^ using three tissue sections stained with H&E and immunohistochemically for p53 and Ki67 (**Fig. 1B**). Only samples for which full agreement was reached were included in our study, and those that received chemotherapy before resection were excluded. Most patients were diagnosed at an advanced stage (>70% stage T3c, **Fig. 1C**), characteristic of sporadic HGSOC ^2^, and had a mean age at diagnosis of 63 years (**Suppl. Table 1**). Importantly, all but one patient had associated invasive carcinoma (IC), which in most cases was sampled in the adnexal region on the same histological slide (**Fig. S1A**), allowing us to directly compare STICs with concurrent ICs. We used additional serial tissue sections to analyze homologous repair deficiency (HRD) in ICs using targeted next-generation sequencing. The HRD status was successfully determined in 29 samples, which classified 15 tumors (approx. half of our cohort) as HRD-positive and 14 tumors as HRD-negative, in excellent agreement with a previous large-scale study conducted by TCGA ^2^. Of the HRD-positive samples, three carried a *BRCA1* mutation and one had a *BRCA2* mutation, classifying them as HRD-positive based both on the genomic instability score (GIS) and the *BRCA1/2* mutations. For three other samples the GIS could not be determined because of the low tumor purity. Pathogenic *TP53* mutations, a ubiquitous genetic feature of HGSOC ^2^, were identified in all 32 sequenced samples, with missense and nonsense mutations being the most prevalent (**Fig. 1C**). Having identified a representative cohort of HGSOC with co-occurring STIC precursor lesions and invasive carcinoma, we next investigated the proteomic landscape.

**Fig. 1:**
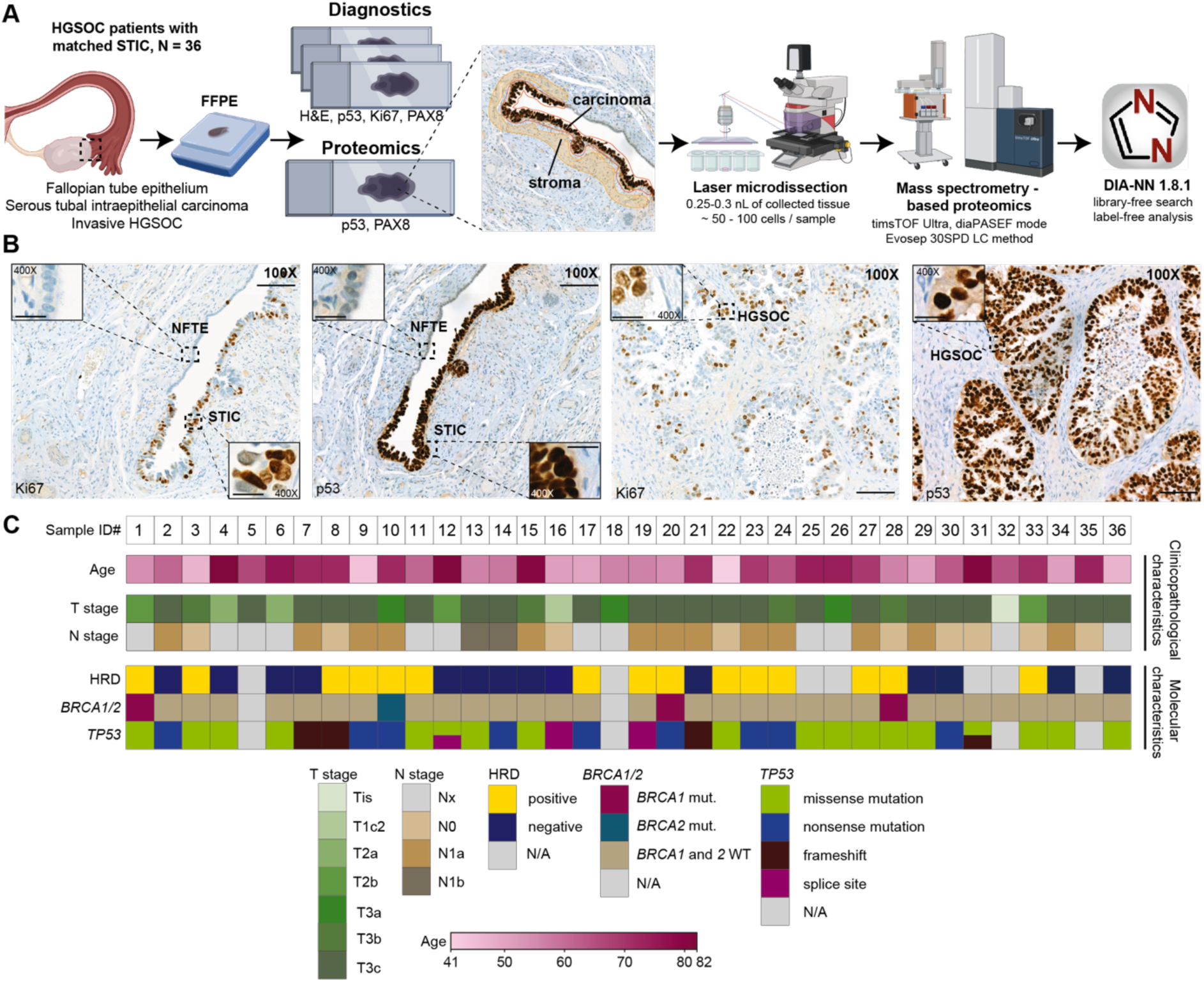
Histology-guided ultra-low input proteomics of HGSOC precursor lesions. **(A)** Pathology-guided ultra-low input proteomics workflow for studying ovarian cancer precursor lesions at the global proteome level. Guided by immunohistochemistry (IHC), healthy and diseased epithelial and stromal regions were laser-microdissected and analyzed using ultrasensitive LC-MS-based proteomics. **(B)** Representative immunohistochemical staining (Ki-67 and p53) of normal fallopian tube epithelium (NFTE), serous tubal intraepithelial carcinoma (STIC), and invasive carcinoma (IC) in the epithelial and stromal compartments. Scale bars: 100X: 100 *μm*, 400X: 20 *μm*. **(C)** Patient cohort (*n* = 36) with clinicopathological and molecular characteristics. Panel (A) was created with BioRender.com

### Deep spatial proteomics quantifies disease-specific alterations at bulk level resolution

We employed our recently developed ultralow-input tissue proteomics workflow ^24^ optimized for the seamless integration of immunohistochemistry (IHC) and immunofluorescence (IF) staining and ultrasensitive LC-MS-based proteomics. This approach enables the profiling of FFPE tissue microregions of only 50-100 cells in size, which is characteristic of STICs. For laser microdissection (LMD), tissues were mounted on PPS metal frame slides and stained with antibodies targeting p53, Ki-67, and PAX8 to guide LMD and proteome profiling. For 35/36 samples, we successfully sampled at least one STIC lesion and one adjacent normal fallopian tube epithelial (NFTE) region. Additionally, in three patients, a second STIC could be sampled from the contralateral fallopian tube. For most cases, a complete set of samples was obtained, including normal epithelial and stromal regions (NFTE and NFT-St), STICs and adjacent stroma (STIC-St), and invasive carcinoma (IC) and its connected stroma (IC-St) (**Fig. 1B and S1A**). IC regions were defined as small tumor ‘nests’ clearly separated from the surrounding stroma (**Fig. 1B**). All samples were processed in a single batch in 384-well low-binding plates using our loss-reduced sample processing protocol (**Methods**) and measured in high-sensitivity diaPASEF mode ^25^ on a timsTOF Ultra mass spectrometer. Raw files were analyzed in DIA-NN using a predicted spectral library, resulting in 10,223 unique protein groups from 192 measurements. The median proteome coverage for epithelial samples exceeded 7,500 protein groups and 6,000 for stromal samples, spanning more than five and four orders of magnitude, respectively (**Fig. 2A-C**, **Fig. S2A-B**). The fewer number of identified proteins in stromal samples can be explained by differences in protein abundance. The top 100 stromal proteins contributed to 37.16% of the total protein mass, compared to 22.96% in epithelial samples. Epithelial and stromal proteomes exhibited high compartment specificity. For example, known HGSOC markers, such as p53, PAX8, and MUC16 (CA125), were specific to the epithelial samples, whereas stromal and immune markers (i.e., collagens, vimentin, CD3E, and CD20) were characteristic of the stromal sample groups (**Fig. 2B-C**, **Fig. S2A-B**). Notably, our high proteome coverage from FFPE tissue regions of only 50-100 cells in size was on par with recent bulk-level studies that required hundreds of micrograms to milligram of fresh frozen tissue ^7,26^. To investigate this further, we compared our dataset with those two large-scale bulk proteomic studies focusing on advanced HGSOC. Comparing of our dataset with that of Qian *et al*. and Zhang *et al*. (CPTAC), we found that 73% of all identified proteins were common in all three studies. Interestingly, our study featured the highest number of uniquely identified proteins (1,022) (**Fig. 2D**), which we attributed to our laser microdissection-based sampling strategy that results in a ‘biological fractionation’ effect for improved proteome coverage. Proteins unique to our dataset were enriched for tumorigenesis-associated processes, such as immunity (interleukin-36 pathway), transcription (RNA polymerase I), and chromatin-related functions (PRC2 chromatin regulator complex, **Fig. 2E**). The biological richness of our proteomics data was also reflected in the hallmark gene sets, which showed a median pathway coverage of 73% (**Fig. 2F**). For example, pathways such as fatty acid metabolism (86%), DNA repair (88%) and oxidative phosphorylation (91%) were almost completely covered by the quantified proteins.

**Fig. 2:**
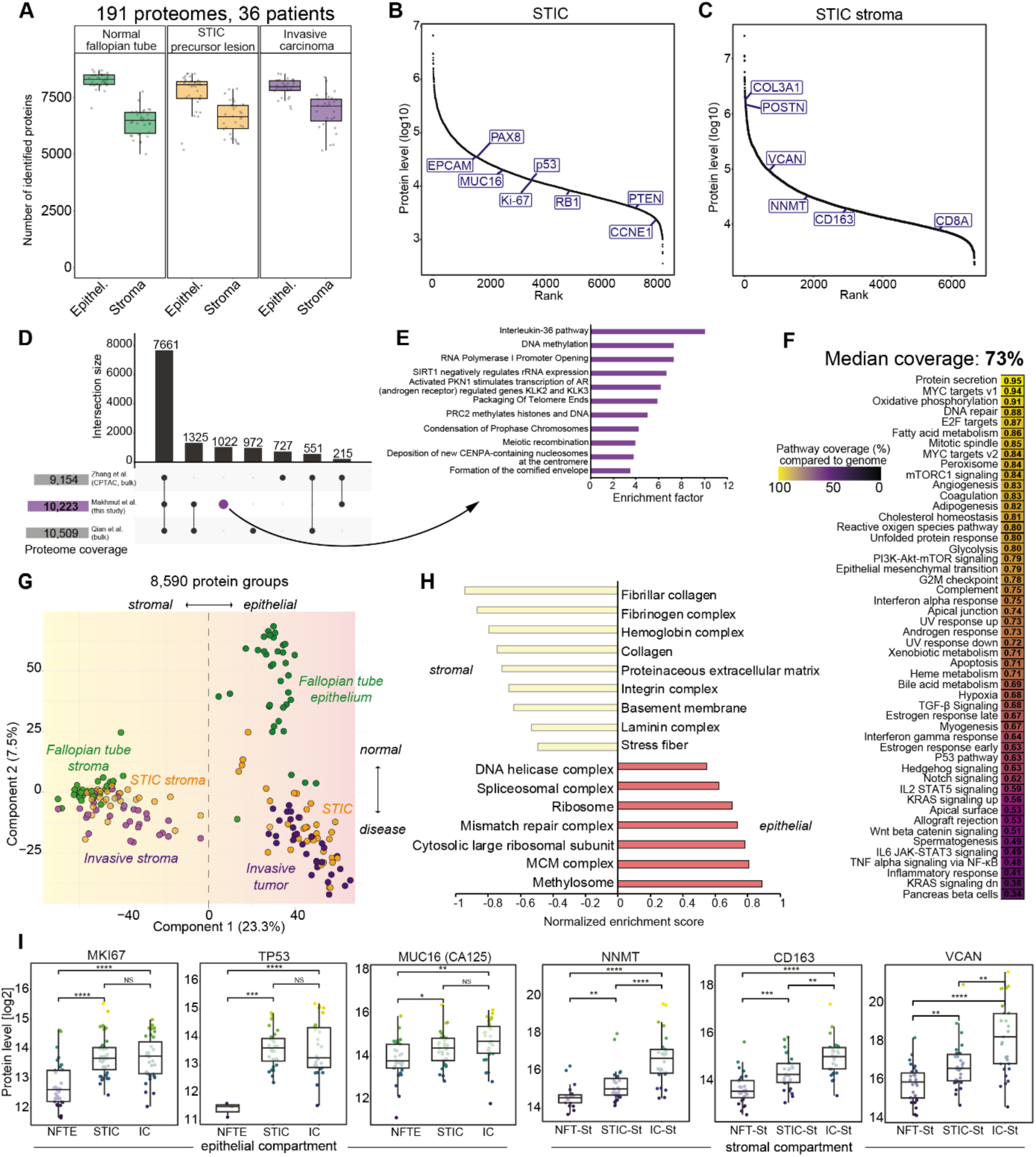
Deep spatial proteomics quantifies disease-specific alterations at bulk level resolution. **(A)** Box plots showing the number of proteins identified in the epithelial and stromal sample groups. Boxplots define the range of the data (whiskers), 25th and 75th percentiles (box), and medians (solid line). **(B)** and **(C)** Dynamic range plots of median protein abundance for epithelial (B) and stromal (C) STIC samples. Known ovarian cancer, cell type, and stromal markers are highlighted. Proteins with 50% valid values for each group are shown. **(D)** Upset plot comparing common and unique proteins identified in this study and in Qian et al.^26^ and Zhang et al.^7^ bulk proteome datasets. **(E)** Overrepresented pathways (Reactome) for the 1,022 proteins uniquely identified in this study (Benjamini-Hochberg FDR < 0.05). **(F)** Median pathway coverage of hallmark gene sets based on the 10,223 proteins identified in this study. **(G)** Principal component analysis (PCA) of all 191 samples based on 8,590 protein groups. **(H)** Pathway enrichment analysis (gene ontology terms) for PC1, separating all epithelial and stromal samples. Representative terms are shown with a Benjamini-Hochberg FDR < 0.05. **(I)** Box plots of the relative protein levels (log2) of the selected epithelial and stromal HGSOC markers. Boxplots define the range of the data (whiskers), 25th and 75th percentiles (box), and medians (solid line). Asterisks indicate two-sided t-test p values (p > 0.05, NS; p < 0.05, *; p < 0.01, **; p < 0.001, ***; p < 0.0001, ****. NFT-St, normal fallopian tube stroma; STIC-St, STIC stroma; IC-St, invasive stroma).

Principal component analysis (PCA) of all 192 proteomes clearly delineated disease-associated changes in the epithelial and stromal compartments, separating healthy control regions from the STIC precursor and invasive regions (**Fig. 2G**). Principal component 1 (PC1, 23.3% total variability) separated epithelial cells from stromal proteomes (**Fig. 2H**), whereas PC2 displayed a transition from healthy to invasive regions in both compartments (7.5% total variability; **Fig. 2G and Fig. S2C**). Dysregulated processes along the progression from healthy to invasive regions included known HGSOC-associated processes, such as increased DNA replication, DNA repair, and inflammation signatures (**Fig. S2C**). Known disease drivers and functional markers are upregulated with disease progression. For example, STICs and ICs were characterized by significantly higher Ki-67 levels, reflecting their high proliferative state, and strongly elevated p53 levels compared with NFTE (**Fig. 2I, p < 0.001**). Interestingly, the strong increase in p53 protein levels was not only specific to tumors with stabilizing missense mutations (compared to p53-null tumors, **Fig. S2D**), but also marked the only significantly expressed protein between these two sample groups, showing that the global proteomes of p53- null and p53-missense tumors were highly related (**Fig. S2D**). The stromal compartment featured strong extracellular matrix and cell-type-related changes associated with disease progression. For example, the cancer-associated fibroblast regulator NNMT ^22^, CD163, a marker of tumor-associated macrophages (TAMs) ^27^, and oncogenic proteoglycan VCAN ^19^ were significantly upregulated from the normal stroma to the invasive stroma (**Fig. 2I**).

Taken together, our deep and compartment-resolved proteomic dataset provides a rich resource for studying the early steps of HGSOC development and progression by incorporating tissue- matched healthy control regions, STIC precursor lesions, and IC.

### STICs and invasive tumors feature strong cell-of-origin signatures

Although there is accumulating evidence that HGSOC originates from fallopian tube epithelial cells ^28,29^, a global and cell-type-resolved proteomic comparison of these putative precursor cells with concurrent STICs and ICs remains elusive. Furthermore, little is known about the proteomic landscape and heterogeneity of STICs and their molecular deviation from NFTE to ultimately form invasive tumors. To address this, we applied a modified Deep Visual Proteomics (DVP) ^30^ workflow to drastically reduce the number of LMD contours required for our cohort, which was distributed over 200 microscopy slides. We isolated small stretches of NFTE guided by PAX8 IHC and isolated concurrent STIC regions and IC ‘nests’ of the same area (total 50,000 µm^3^). Principal component analysis revealed strong proteome differences between the three sample groups, with NFTE clearly grouping apart from the disease states (**Fig. 3A**). Interestingly, four NFTE samples grouped much closer to STICs and ICs, indicating that they likely did not represent normal regions (here referred to as premalignant). Proteins upregulated in these samples were strongly enriched for cancer-associated pathways, such as higher DNA replication, cell cycle, and DNA double-strand break repair signatures (**Fig. 3B, D**). The most upregulated protein was the cell cycle regulator CDKN2A (p16/INK4A, **Suppl. Table 2**), an established histological marker for the diagnosis of STIC with a p53 null phenotype ^31^. Informed by proteomics, we therefore re-stained these tissues against p16 and evaluated whether they were normal-like or indeed malignant lesions. Indeed, we observed strong and uniform p16 staining (**Fig. 3C**) and a morphology suspicious for STIC for two of them, leading to the re-diagnosis of two of these samples as serous tubal intraepithelial lesions (STIL)^32,33^ and STIC for the other two. The absence of p53 positivity (p53 null phenotype) and only marginal morphological changes in the epithelium potentially led to their misclassification. We also noticed that two samples initially labeled as STIC grouped closer to the NFTE. Reanalysis of these samples revealed that one was the only STIC sample with no associated IC, likely representing an earlier disease state captured by proteomics. The second one represented an STIC with a stabilizing p53 missense mutation but a p53 null phenotype in the invasive component, suggesting an incidental STIC scenario. These observations clearly underscore the potential of our spatial proteomics approach as a companion diagnostic tool. Pairwise comparison between the four premalignant fallopian tube lesions and the normal epithelium also revealed the presence of ciliated cell signatures in the NFTE samples (**Fig. 3B, D**), potentially because of the co-isolation of some ciliated cells. To investigate this further, we integrated data from a recent single-cell transcriptomic atlas of the healthy human fallopian tube ^34^ to assess which cell type and cell states were present in our dataset. This resulted in binary clustering, marking roughly half of our NFTE samples as secretory cell-enriched and the other half as more ciliated (**Fig. S3B, Suppl. Table 4)**. As expected, the cell type marker PAX8 (secretory cells), expressed in 96% of HGSOCs ^35^, and the transcription factor FOXJ1 (ciliated cells) were among the most significantly regulated proteins in these two clusters (**Fig. 3E, Suppl. Table 5**). Motivated by this finding, we next asked whether these two distinct cell-type signatures present in our dataset could help to unmask cell-of-origin signatures preserved in STICs and invasive tumors. We found that the secretory-like NFTE cluster grouped closely with all malignant stages (STIC and IC) and was clearly apart from the ciliated-like cluster (**Fig. 3F**). Co-expressed markers in the carcinoma/secretory NFTE group included known histological markers, such as PAX8 and STMN1, a proposed ancillary marker for detecting STIC lesions with a p53 null phenotype ^31^. Other proteins with similar cell-type specific and disease-associated expression included DHCR24, an oxidoreductase important in cholesterol biosynthesis ^36^, the collagen-specific chaperone SERPINH1, the calcium binding protein S100A4, as well as the basal cell adhesion protein BCAM, as well as many more (**Fig. S3D**). To further validate that STICs are globally related to normal secretory cells, we performed additional immunofluorescence whole-slide imaging of six tissue sections and precisely isolated PAX8+ and PAX8- cells for direct proteomic comparison (**Fig. 3H-I, Fig. S3E-G**). PAX8+ secretory cell proteomes showed strong overlapping signatures with STICs and clustered apart from all PAX8- epithelial samples (**Fig. 3J-K and Fig. S3H**). Secretory cell-specific signatures were strongly enriched in the STIC/PAX8+ cluster with high p16, p53, and PAX8 protein levels and pathways related to cell cycle, unfolded protein response and DNA repair. In contrast, PAX8- cells expressed high levels of ciliated-cell-specific cilium and intra- flagellar transport proteins, as well as FOXJ1 (**Fig.3K, Fig. S3F-I, Suppl. Table 6-9**). The normal epithelial cluster, which comprised PAX8+ and PAX8- cells, was enriched for previously identified transition signatures between secretory and ciliated cells (unclassified clusters 2 and 3) ^34^, consistent with the expected tissue composition and possibly capturing the previously proposed secretory-to-ciliated cell differentiation programs in the fallopian tube epithelium.

**Fig. 3:**
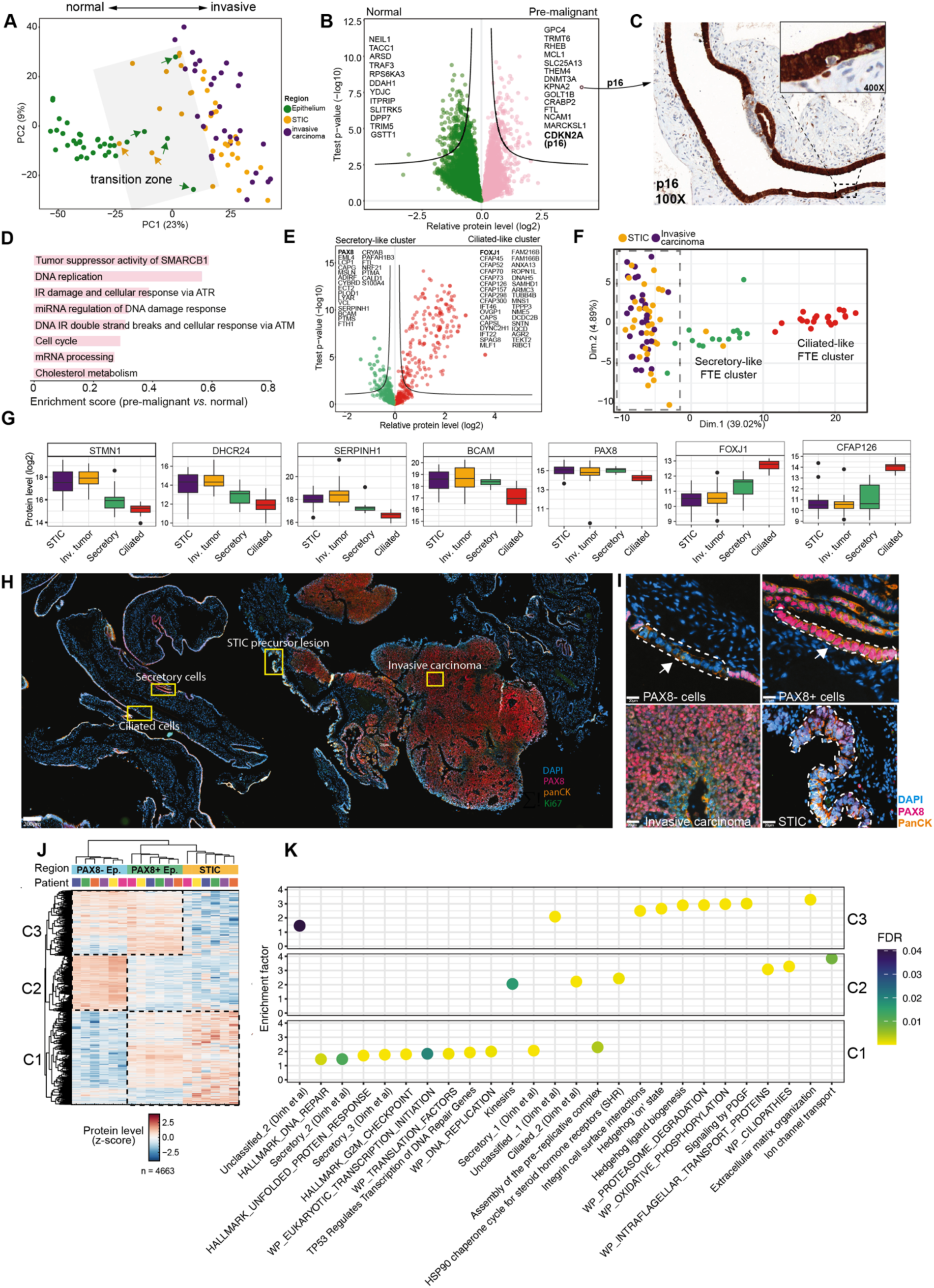
STIC and invasive carcinomas feature strong cell-of-origin signatures. **(A)** Principal component analysis (PCA) of 101 normal epithelium, STIC, and invasive carcinoma proteomes based on ANOVA-significant 3564 proteins. **(B)** Volcano plot of pairwise proteomic comparison between pre-malignant (light-pink) and normal fallopian tube epithelium (green) samples. Markers with the highest fold change are highlighted (two-sided t-test, permutation-based FDR < 0.05). **(C)** Representative immunohistochemical staining of CDKN2A (p16) in one pre-malignant fallopian tube epithelial region. **(D)** Pathway enrichment analysis based on the t-test difference between premalignant and normal fallopian tube epithelial samples. Selected significantly enriched pathways are shown for pathways higher in pre-malignant samples (Benjamin-Hochberg FDR < 0.05) **(E)** Volcano plot of the pairwise proteomic comparison between ciliated-like epithelial (red) and secretory-like epithelial (green) samples. Known secretory (e.g., PAX8) and ciliated (e.g., FOXJ1) markers are highlighted (two-sided t-test, permutation-based FDR < 0.05). **(F)** Principal component analysis of all epithelial samples based on 632 proteins overlapping with cell type-specific markers identified by single-cell transcriptomics^34^. **(G)** Boxplots of the relative protein levels (log2) in panels **(G)** and **(H)**. Boxplots define the range of the data (whiskers), 25th and 75th percentiles (box), and medians (solid line). **(H)** Immunofluorescence image of one representative HGSOC tissue section stained for PAX8, Ki-67, panCK, and DAPI (DNA). Yellow boxes highlight regions of invasive carcinoma, STIC precursor lesion, and normal fallopian tube ciliated and secretory cells, which were sampled for proteomic profiling). **(I)** Magnified tissue regions from immunofluorescence images in panel **(H)**. Arrows show exemplary epithelial regions of the PAX8+ and PAX8-regions used for proteomic profiling. Scale bar = 20 *µm*. **(J)** Unsupervised hierarchical clustering of all ANOVA significant proteins between STICs, secretory, and ciliated fallopian tube epithelial cells obtained from six patients. Relative protein levels (z-scores) are shown. Three distinct clusters, C1, C2, and C3, are highlighted. PAX8+ secretory cells clustered together with STICs and were separated from ciliated PAX8− samples. **(K)** Pathway enrichment analysis (Reactome, Hallmark, WikiPathways, and cell type signatures from Dinh *et al.* ^34^) showed significantly overrepresented pathways for clusters C1, C2, and C3 with a Benjamini-Hochberg FDR < 0.05.

In summary, this cell-type resolved proteomic analysis revealed strong cell-of-origin signatures present in STIC and concurrent carcinomas, supporting the view that HGSOC originates in secretory cells of the distal FT. Moreover, our analysis highlighted several new proteins as potential ancillary markers for the detection of p53 null phenotype STIL and STIC lesions, independent of HRD and *TP53* mutation status.

### High proteome similarity between STICs and concurrent invasive carcinomas is reflected in distinct tumor-immune phenotypes

Next, we compared all STICs and ICs. Similar to principal component analysis (PCA) (**Fig. 3A**), unsupervised hierarchical clustering of the 2,000 most variably expressed proteins in our dataset confirmed that STICs and ICs were highly related, resulting in two distinct clusters, irrespective of sample type (STIC or IC) (**Fig. 4A-B and S4A**). Clustering was also independent of *TP53* mutation type, histological stage, and HRD status, but showed an association of cluster 2 samples with higher patient age (**Fig. 4A and Fig. S4B**). Nearly all (29 / 30) patient-matched STIC-IC pairs clustered together (**Fig. 4A-B)**, underlining the strong patient-specific proteome profiles. The high similarity between tissue-matched STICs and ICs was also reflected in their higher global proteome correlation (median Pearson *r* = 0.94, **Fig. S4C**) compared to inter-patient comparisons (median Pearson *r* = 0.91). Interestingly, bilateral STICs obtained from both ovaries of the same patient featured exceptionally high proteome correlations (Pearson *r* = 0.97), despite being spatially unrelated. While this pointed towards a possible clonal relatedness of these two premalignant lesions, additional genetic analyses are needed to confirm or reject this hypothesis.

**Fig. 4:**
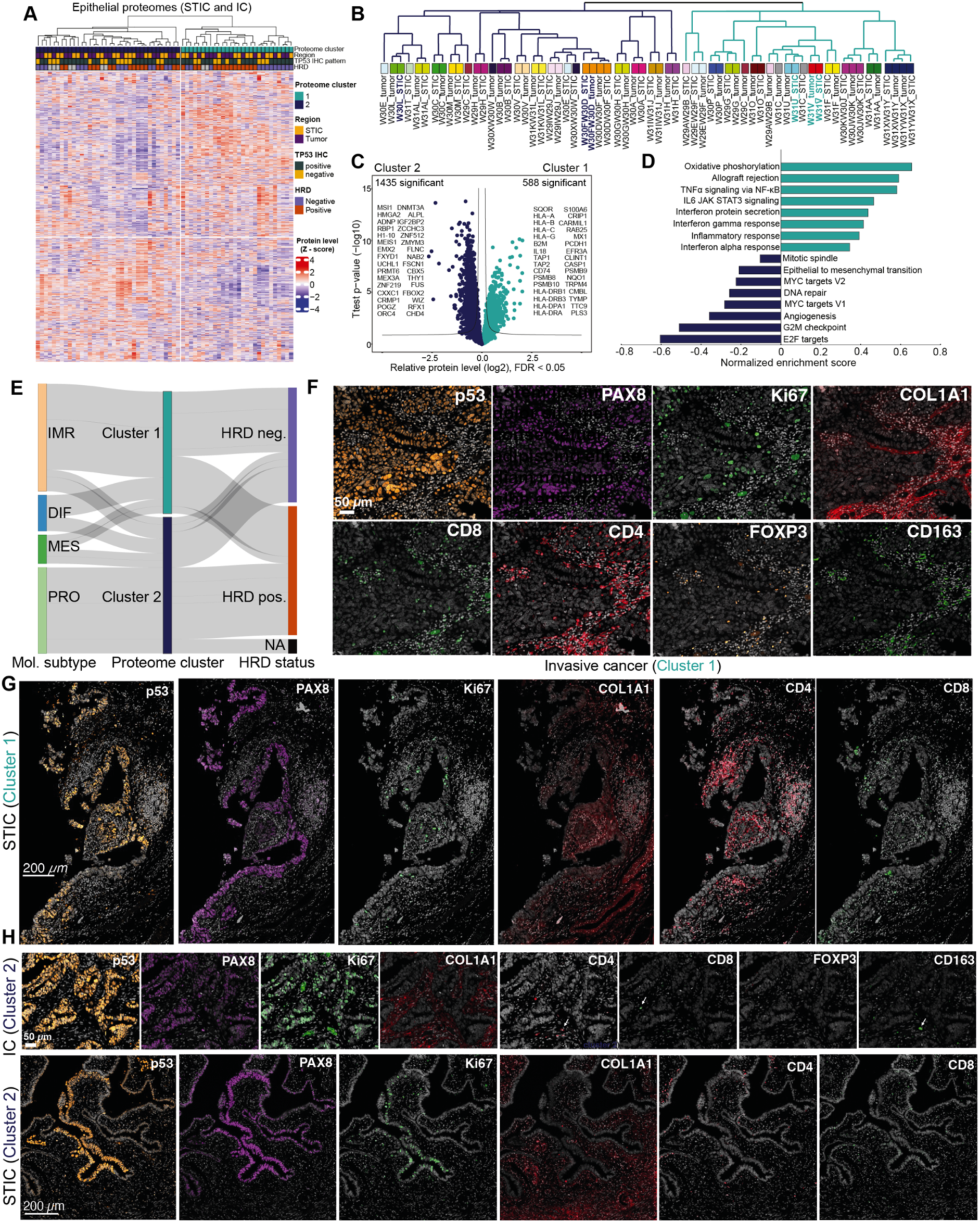
High proteome similarity between STICs and IC and identification of commonly dysregulated pathways. **(A)** Unsupervised hierarchical clustering of all 64 STIC and IC samples based on the 2,000 most variable proteins (highest median absolute deviation), showing two clusters. Relative protein levels (z-score) are shown with clinicopathological information as color bars. **(B)** Cluster dendrogram related to panel (A). Selected cluster 1 and 2 patients shown in panels 4F-H are highlighted in bold. **(C)** Volcano plot of pairwise proteomic comparison between cluster 1 (turquoise) and cluster 2 (dark blue) samples. Proteins with the highest fold change are highlighted (two-sided Student’s t-test, false discovery rate [FDR] < 0.05). **(D)** Pathway enrichment analysis (WikiPathways, Hallmarks) based on the t-test difference between cluster 1 and cluster 2 epithelial samples. Selected pathways with a Benjamin-Hochberg FDR < 0.05 are shown. **(E)** Sankey plot illustrating the distribution of STIC & IC samples across consensusOV molecular subtypes, cluster 1 or cluster II, and HRD statuses. The width of each flow corresponds to the proportion of the samples within each category, highlighting the relationship between molecular subtype, cluster assignment, and HRD status. **(F - H)** CyCIF of one representative immune-enriched cluster 1 tumor and STIC (patient W31V, panel F – tumor, panel G - STIC) and one immune-deserted cluster 2 tumor and STIC (patient W30FW30D, panel H, top – tumor, bottom - STIC) confirms pronounced differences in Ki-67+ tumor cells and intra-tumoral immune cell infiltration. Scale bar = 50 *µm* (STIC), 200 *µm* (tumor).

Cluster 1 tumors were primarily enriched for immune-associated processes (e.g., interferon response, TNFα signaling, antigen processing, and presentation), whereas cluster 2 samples showed elevated levels of cell cycle, EMT, and DNA repair associated proteins (**Fig. 4C-D, Suppl. Table 10-11**). To integrate our findings with previously identified HGSOC molecular subtypes ^2,7^, we employed the consensusOV classifier ^37^ and assessed whether known subtypes were present in our two main clusters. 36 proteomes could be assigned to one subtype based on a margin score of greater than 0.2 (difference between the probabilities of most-probable and second most-probable subtype, mean score = 0.54). Notably, most cluster 1 samples (12 of 17, 71%) were classified as the ‘immunoreactive’ (IMR) subtype, whereas cluster two samples were classified as ‘proliferative’ (PRO) tumors, independent of HRD status (**Fig. 4E**). Only a few samples were classified as mesenchymal (*n*=4) or differentiated (*n*=5). We mainly attribute this discrepancy to our compartment-resolved strategy that contains minimal stromal admixing, thereby enabling more accurate tumor subtyping. Using single-cell RNA sequencing, previous studies have contextualized HGSOC bulk subtypes and identified differentiated (DIF) and proliferative (PRO) subtypes as tumor cell-specific signatures ^4^. In contrast, stromal and immune cells are associated with the mesenchymal and immunoreactive subtypes, respectively^38^. The study by Geistlinger *et al*. also revealed that the DIF subtype is associated with the strongest lymphocyte infiltration, letting us speculate that cluster 1 tumors could likely represent DIF tumors whose proteomes were strongly masked by pronounced intra-tumoral immune infiltration. Indeed, not only was the second-best subtype assignment for IMR tumors the DIF subtype (**Fig. S4G**), but cyclic immunofluorescence imaging (CyCIF) also confirmed strong immune cell infiltration in this subgroup (CD8+ cytotoxic T cells, CD4+ helper T cells, FOXP3+ regulatory T cells, CD163+ M2-like TAMs, and CD11c+ dendritic cells) (**Fig. 4F-G**). Conversely, cluster 2 tumors showed negligible immune infiltration based on proteomics and CyCIF, but had a larger fraction of Ki-67+ tumor cells, in line with their proliferative nature (**Fig. 4H**).

We next addressed whether STICs and ICs featured similar molecular subtypes and whether the progression from STIC to IC was associated with a change in molecular subtype identity. Ten matching STIC-IC pairs could be clearly assigned to one of the four subtypes. We found a similar subtype frequency at both stages (**Fig. S4F**), suggesting a similar level of phenotypic and proteomic heterogeneity. For the majority of paired cases (70%), the STIC and IC subtypes were coherent (**Fig. S4H**), underlining the absence of a distinct epithelial STIC proteotype distinguishable from ICs. This finding was supported by the global proteome comparison, which revealed only a few differentially regulated proteins (9 out of 6,744) between all STIC and IC samples (**Fig. S4I**) and exceptionally high proteome correlations of patient-matched STIC-IC pairs (**Fig. S4C**).

#### Progressive ECM remodeling and co-evolution of an immunosuppressive tumor microenvironment

We next focused our attention on the tumor microenvironment (TME). The TME comprises various stromal and immune cell types and the extracellular matrix (ECM) and is critically involved in all phases of tumorigenesis^39–41^. As the TME is increasingly recognized as a promising therapeutic target across cancer entities ^42^, quantitative proteomics data comparing normal, precursor and invasive stromal tissue regions are particularly important. While the broad dynamic range of protein abundance in the ECM generally poses a significant obstacle for MS-based protein identification ^43^, our analysis yielded more than 6,000 proteins for each stromal microregion, distributed over four orders of magnitude (**Fig. 2A-B**). Principal component analysis revealed a disease gradient from normal stroma (NFT-St) over STIC stroma (STIC-St) to invasive stroma (IC-St) (**Fig. 5A**). Interestingly, in contrast to our epithelial findings that showed that STICs and ICs were proteomically highly related and distinct from the normal epithelium, STIC-St proteomes showed both normal and malignant signatures. This observation was supported by unsupervised hierarchical clustering of the 2,888 differentially abundant proteins, representing roughly one-third of our analyzed stromal proteome (**Fig. 5B, Suppl. Table 12**). The invasive stroma cluster showed high inflammation, hypoxia and VEGF signaling accompanied with strong lymphocyte and myeloid cell signatures (e.g., T-cell, B-cell, macrophage, and NK-cell). Conversely, the normal-like cluster, which included half of the STIC stroma group, was enriched for fibroblast and endothelial cell signatures, as well as core ECM functions, indicating a structurally different and more intact ECM. This prompted us to more systematically analyze the quantitative ECM-changes during HGSOC development and to this end filtered our data for all quantified matrisome-associated proteins, as recently catalogued ^44^. From the total of 467 matrisome related proteins in our dataset, we identified 95 (20%) as differentially abundant between normal and invasive stroma (**Fig. 5C, Suppl. Table 13**). Proteins with the highest significance included known ECM degraders, such as cathepsins (CTSA, CTSC, CTSS, and CTSZ), matrix metalloproteinases (MMP11 and MMP14), pro-inflammatory cytokines (TGFϕ31, TGFϕ32, IL16, and IL18), and insulin-like growth factor-binding proteins (IGFBP2, IGFBP7). We identified strong structural ECM changes, as evident from the downregulation of multiple collagen isoforms (e.g., collagen types 4, 6, 11, and 21) during HGSOC progression (**Fig. 5C**). Our collagen isoform-resolved data also enabled us to quantify a common decrease in the COL3/COL1 ratio in 27 of 32 tissues (84% of the cohort), a marker for fibrosis, ECM stiffening, and tumor progression ^45,46^ (**Suppl. Table 14**). An exception to the general decrease in collagen abundance with disease progression were collagens 8A1, 8A2, and 10A1, which showed higher levels in the invasive stroma (**Fig. 5D**). Notably, these three collagen isoforms were previously linked to myofibroblasts (myCAFs) in pancreatic cancer and were associated with unfavorable clinical outcomes ^47^. This underlines the central role of CAFs as key drivers for oncogenic ECM remodeling in our cohort, which was further supported by a consistent up-regulation of several other CAF markers, such as NNMT, TNC and FAP (**Suppl. Table 9**). We also identified other collagen isoforms, such as COL9A3, COL9A1, COL23A1, and COL13A1, not previously reported in HGSOC and whose specific functions remain to be elucidated.

**Fig. 5:**
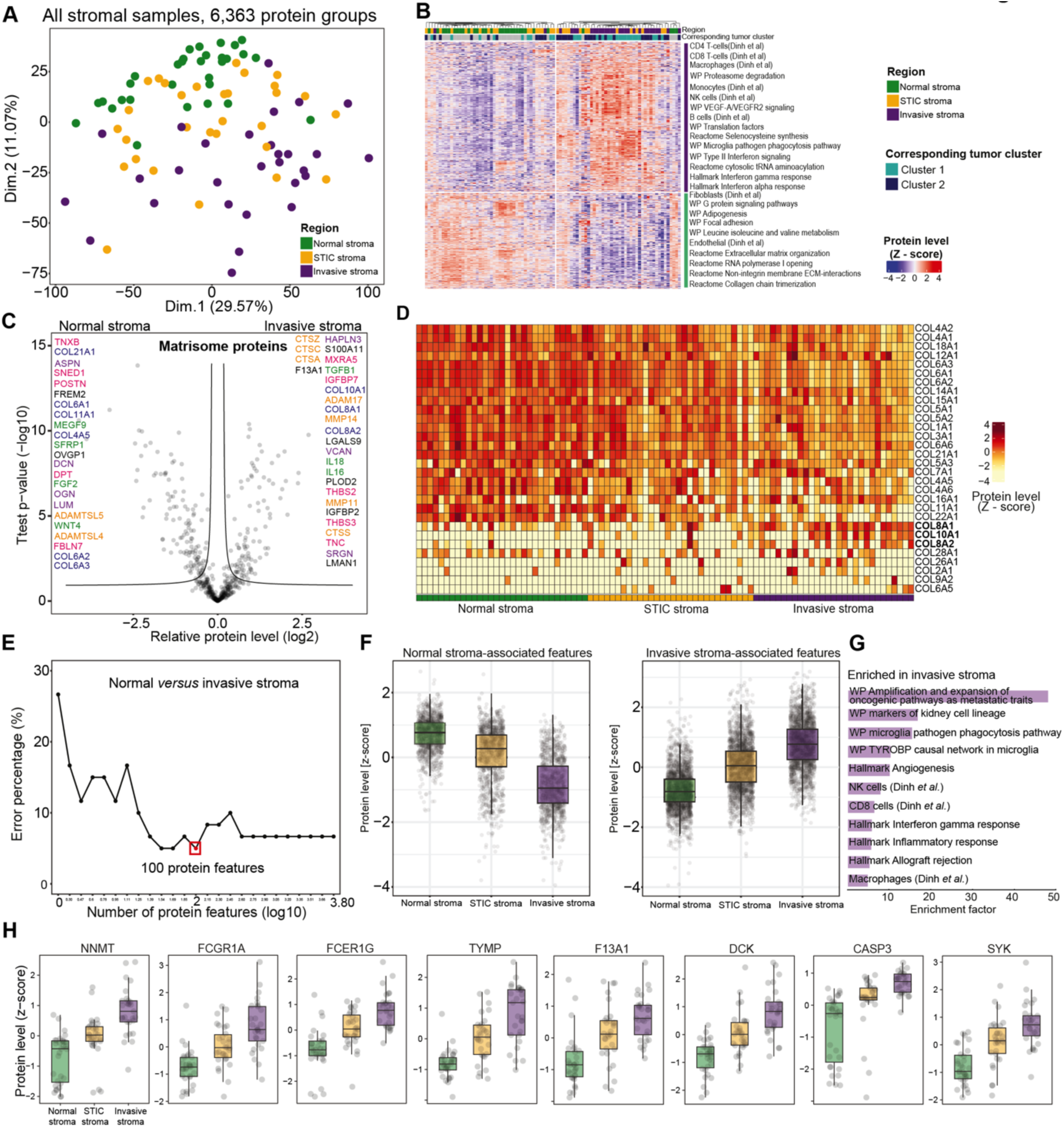
Quantitative remodeling of the tumor microenvironment during disease progression. **(A)** PCA of all 90 stromal proteomes based on 6,363 protein groups. PC1 and PC2 accounted for 40.64% of the total data variability. **(B)** Unsupervised hierarchical clustering of all stromal proteomes based on 2888 ANOVA-significant proteins (permutation-based FDR < 0.05). Relative protein levels (z-score) are shown with clinicopathological and tumor subtype information as color bars. A subset of enriched pathways (WikiPathways, cell-type signatures obtained from Dinh *et al.*^34^) for the two main row clusters marked in purple and green is shown. **(C)** Pairwise proteomic comparison between invasive and normal stroma (two-sided t-test) for all matrisome-associated proteins. Proteins with a permutation-based FDR < 0.05 were considered significant. Orange: ECM regulators; green: secreted factors, pink: ECM glycoproteins; blue: core matrisome; purple: proteoglycans. **(D)** Relative protein levels (z-score) of all quantified collagens across normal, STIC, and invasive stroma. **(E)** Support vector machine-based identification of top-ranked protein features distinguishing normal from invasive stroma. **(F)** Boxplots of the relative protein levels (z-score) of the top 100 proteins distinguishing normal stroma from invasive stroma. Left: Proteins that were higher in the normal stroma. Right: Proteins that were higher in the invasive stroma. **(G)** Overrepresented pathways (FDR< 0.05) of the top 100 proteins compared to all proteins in the dataset. **(H)** Boxplots of relative protein levels (z-score) of FDA-approved drug targets.

To prioritize protein drug targets significantly upregulated in the invasive stroma, we next employed support vector machine (SVM) classification combined with feature ranking and retrieved a 100-protein stromal signature that robustly (5% error rate) separated all normal and malignant stromal proteomes (**Fig. 5E-F**). Notably, protein levels of the STIC stroma samples were in between normal and invasive stroma levels, indicating a gradual decrease or increase of the selected proteins. Our signature included many known proteins of the desmoplastic stroma previously linked to poor patient outcome ^19^ (e.g., VCAN, THBS2 and TNC), proinflammatory cytokines (IL-18), immune response and complement system modulators (e.g., F13A1, FCGR1A and FCER1G) and markers of tumor promoting cell-types (e.g., NNMT for CAFs and CD163 and MSR1 for M2-polarized TAMs)^22,27,48^ (**Fig. 5G and S5B-C, Suppl. Table 15**). Notably, nine proteins upregulated in the invasive stroma (NNMT, FCGR1A, FCER1G, TYMP, F13A1, DCK, CASP3, and SYK) represented FDA-approved drug targets outside of ovarian cancer (**Fig. 5F**), emphasizing their high relevance for future preclinical investigations.

### Identification of early dysregulated pathways of therapeutic relevance

To identify commonly dysregulated pathways during early HGSOC development, we compared all NFTE to STICs. Such data are of high translational importance, for example for the development of new disease prevention and therapeutic strategies ^49^. To this end, we compared all secretory-like NFTE samples with their corresponding STICs. Overall, we observed pronounced proteomic differences with 615 differentially abundant protein groups (**Fig. 6A, Suppl. Table 16**), in stark contrast to the STIC *versus* IC comparison that only yielded nine differentially expressed proteins (**Fig. S4I**). Proteins of highest significance and fold change included DNA replication proteins (e.g., MCM2,3,4,5 and 7), the cell cycle regulator CDKN2A (p16-INK4A), the microtubule and cell cycle regulator STMN1, and the insulin growth factor binding protein IGFBP2, which was previously reported to be upregulated in STICs through DNA hypomethylation ^50^. Notably, we also discovered several dysregulated metabolic pathways (**Fig. 6B-C, Fig. S6A**). For instance, protein levels related to prostaglandin biosynthesis were higher in NFTE than in STIC and IC (**Fig. 6C**), reflecting the important physiological role of the fallopian tube in hormone regulation ^51^, and the loss of normal epithelial cell function during carcinogenesis. Conversely, the glycolysis and cholesterol biosynthesis/mevalonate pathways were prominently elevated in carcinomas, coinciding with a decrease in oxidative phosphorylation (OxPhos). Our data further showed that these metabolic pathway changes were independent of the molecular subtype (**Fig. 6D and Fig. S6B, Suppl. Table 18**), suggesting that metabolic adaptation to a glycolytic and cholesterol-dependent state is a common and likely early event during HGSOC development. We found several key enzymes of the cholesterol biosynthesis pathway were up-regulated in STICs and ICs compared to normal epithelial cells (**Fig. 6E**). Most prominently, the enzymes dehydrocholesterol-reductase 24 (DHCR24) and 7 (DHCR7) were up-regulated by 3.3 and 5.2- fold, respectively, which catalyze the final steps in cholesterol biosynthesis. Integrating a large-scale pan-cancer study ^52,53^ comparing 1,739 cell lines of different tumor origin confirmed high DHCR24 and DHCR7 mRNA levels in HGSOC (**Fig. S6C-D**). Next, we asked if blocking the cholesterol biosynthesis pathway via dehydrocholesterol-reductases could offer a novel approach for HGSOC treatment. To test this, we treated four ovarian cancer cell lines (OAW- 42, OVCAR-8, ES-2, and EFO-21) of high molecular similarity to HGSOC tumors ^54^ with the DHCR7 inhibitor AY-9944. For comparison, we included the cholesterol inhibitor simvastatin and four doses of carboplatin, the main chemotherapeutic for HGSOC treatment. On average, DHCR7 inhibition at 10 µM showed a significant 67% reduction in cell viability, while simvastatin was effective in only two of the tested cell lines (OVCAR8 and ES-2), which were also most sensitive to AY-9944 (**Fig. 6F**). Carboplatin treatment resulted in variable responses consistent with the expected levels of resistance. Notably, we observed significant drug synergy between the DHCR7 inhibitor and carboplatin based on three reference models (mean HSA score = 15.24, *p* = 1.65E-38; mean Bliss score = 10.46, *p* = 5.22E-18; mean ZIP score = 10.31, *p* = 2.03E-19), as evident from OVCAR-8 cells treated at various drug concentrations (**Fig. 6G and Fig. S6E-H**).

**Fig. 6:**
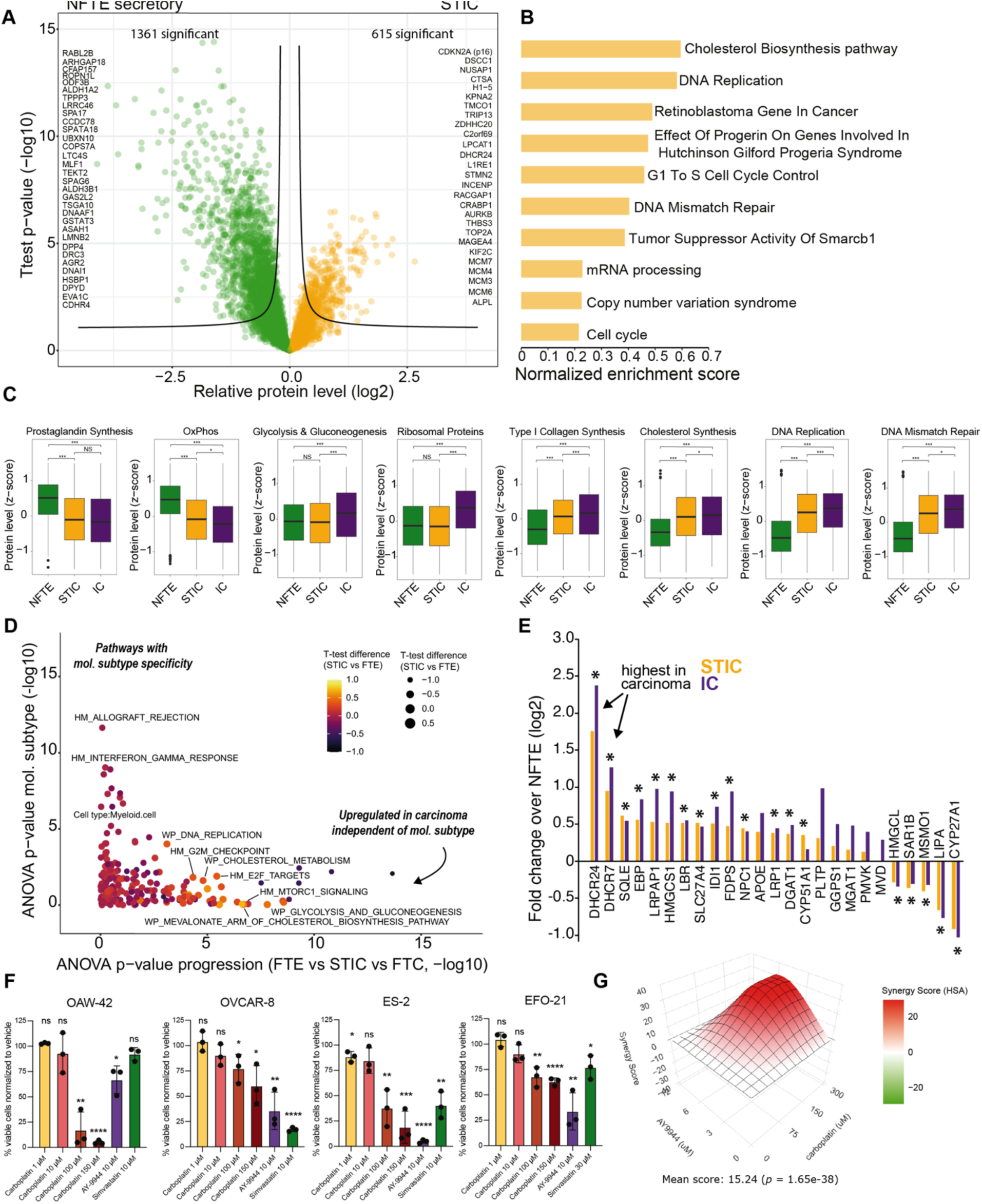
Identification of commonly dysregulated pathways of therapeutic relevance. **(A)** Volcano plot of the pairwise proteomic comparison between STIC (yellow) and secretory-cell-enriched NFTE samples (green). Proteins with the highest fold change were highlighted (two-sided t-test, permutation-based FDR < 0.05). **(B)** Pathway enrichment analysis (WikiPathways, Benjamin-Hochberg FDR < 0.05) revealed processes upregulated in STICs. **(C)** Boxplots of relative protein levels (group average, z-score) for selected pathways. Boxplots define the range of the data (whiskers), 25th and 75th percentiles (box), and medians (solid line). OxPhos: oxidative phosphorylation. **(D)** Two-dimensional ANOVA showing pathways significantly affected by disease progression (x-axis) and molecular subtype (y-axis). Note that metabolism-related terms such as the mevalonate and cholesterol biosynthesis pathways are upregulated in carcinoma, independent of molecular subtype. **(E)** Ranked protein fold changes (STIC and IC *vs.* NFTE, log2) of all quantified cholesterol biosynthesis/metabolism pathway proteins. Most proteins show higher expression in STICs and ICs. Proteins significantly changed in both comparisons (STIC *vs.* NFTE and IC *vs.* NFTE, FDR < 0.05) are highlighted with asterisks. **(F)** Proliferation assay of ovarian cancer cell lines (OAW-42, OVCAR-8, ES-2, and EFO-21) treated with different doses of carboplatin, the DHCR-7 inhibitor AY-9944 and simvastatin. Treatment effect relative to matched vehicle. Experiments were performed in biological triplicates (with n = 3 technical replicates each). Mean ± SD; significance test: unpaired Student’s t-test: ns (not significant), ∗p < 0.05, ∗∗p < 0.01, ∗∗∗p < 0.001, ∗∗∗∗p < 0.0001. **(G)** Synergy score map of OVCAR-8 cells treated with different concentrations of carboplatin and the DHCR7 inhibitor AY9944. The mean HAS (Highest Single Agent) score was 15.24 (p = 1.65 × 10^−38^), showing a statistically significant synergistic effect of the two tested drugs.

Together, our patient and *in vitro* data support the notion that blocking the cholesterol biosynthesis pathway, possibly through dehydrocholesterol-reductase inhibition, could be a promising and previously hidden avenue for ovarian cancer treatment.

In summary, we identified commonly dysregulated pathways linked to early HGSOC pathogenesis, thus providing a rich resource for future biomarker research and drug target discovery.

## Discussion

Despite the common view that STICs represent the precursor lesion for the majority of HGSOCs, their global proteomic landscape and relationship with normal epithelium and concurrent invasive tumors have remained largely unaddressed. Building on our previously developed frameworks for deep spatial proteomics of ultra-low-input tissue ^24^ and deep visual proteomics ^30^, we performed a comprehensive proteomic analysis of HGSOC precursor lesions quantifying over 10,000 proteins. Such deep proteome coverage was previously obtained from hundreds of micrograms to milligrams of freshly frozen tissue ^7,26^. Closely guided by histopathology and whole-slide imaging, this approach enabled us to dissect early changes in the disease-related proteome and to unravel the proteomic heterogeneity of STICs. We quantitatively compared the proteome of STICs and adjacent normal fallopian tube epithelial cells, the assumed cell of origin for HGSOC, and revealed hundreds of disease-related protein level changes. Compared to traditional bulk approaches using tumor-adjacent control tissue for comparison, mostly comprised of stromal cells and ECM, our cell-type resolved approach is more accurate in identifying disease-specific alterations and functional drivers. We provide strong support for the secretory fallopian tube epithelial cell as the cell-of-origin for our analyzed tumors. Our paired proteome analysis of STICs and adjacent healthy fallopian tube epithelial cells revealed that STICs and PAX8+ epithelial cells were not only globally highly related, compared to PAX8- epithelial cells, but also co-expressed well-established histological HGSOC and secretory cell markers (e.g., PAX8, STMN1), as well as several newly identified ones (e.g., DHCR24, BCAM, and SERPINH1). These findings are in line with recent single-cell sequencing data ^34,55^ as well as HGSOC mouse models ^14,29^, supporting the view that the majority of HGSOC originates from secretory cells of the distal fallopian tube.

Surprisingly, we uncovered four pre-malignant tissue regions that were missed by histopathological examination, likely due to p53 staining negativity (p53 null phenotype) and ambiguous Ki-67 status. However, our deep proteome data clearly marked these regions as pre- malignant with a strong upregulation of DNA replication, DNA damage response, and cell cycle proteins, including CDKN2A (p16), which is an additional marker used to diagnose STILs and STICs ^31^. Morphological and immunohistochemical reevaluation by p16 confirmed our suspicion and allowed us to reclassify them as pre-malignant. In accordance with the current pathological nomenclature, these regions were serous tubal intraepithelial lesions (STILs). Unbiased spatial proteomics data hence show great potential for companion diagnostics, especially in cases difficult to diagnose by morphology and IHC alone, or when lacking disease-specific histological markers

Many of the identified proteins upregulated in STICs and invasive carcinoma have known roles in cell cycle regulation, DNA metabolism, and oncogenic signaling, characteristic of this highly proliferative and genetically unstable cancer. We found that metabolic reprogramming is a common and likely early event in HGSOC pathogenesis, identifying several onco-metabolic changes for therapeutic intervention. For example, our data revealed that oxidative glucose metabolism gradually decreases from healthy fallopian tube epithelial cells, over STICs, to invasive tumors, accompanied by upregulated glycolysis and lipid metabolism (cholesterol biosynthesis/mevalonate pathways). While the metabolic switch towards a glycolytic state is a well-known characteristic of many cancers ^56^, upregulation of cholesterol biosynthesis is intriguing. Studies in breast and liver cancer showed a link between *TP53* mutant tumors and increased cholesterol biosynthesis ^57,58^, likely explaining this signature’s dominance in our *TP53* mutant cohort. Importantly, this metabolic phenotype was independent of molecular subtype, supporting the view that blocking cholesterol biosynthesis ^59^, possibly via the up- regulated enzymes DHCR7 or DHCR24, could be a promising therapeutic strategy for high-grade serous ovarian cancer. Our *in vitro* data based on several bona-fide HGSOC models established that inhibiting the terminal steps of cholesterol biosynthesis via DHCR7 inhibition not only shows dose-dependent tumor cell killing, but also synergizes with platinum-based chemotherapy. Our data suggest a possible metabolic dependency of HGSOC on altered lipid and cholesterol metabolism with prospects for combination treatments to improve therapeutic outcomes and to overcome drug resistance. Notably, a previous study linked the inhibition of the cholesterol synthesis pathway by statins to decreases STIC formation in mouse models of ovarian cancer ^60^, further supporting the relevance our findings. However, more preclinical research is needed to assess our data’s translational relevance in HGSOC therapy.

We further demonstrated that STICs and invasive carcinomas show high proteomic similarity, lacking a distinct STIC proteotype distinguishable from advanced carcinomas. Our results establish that STICs mirror the phenotypic and molecular subtype heterogeneity of ICs, implying they exhibit the full oncogenic potential of invasive HGSOC. Two recent smaller-scale studies ^61^, including our own ^22^, support this view. One explanation is that at HGSOC diagnosis, STICs have likely undergone additional molecular aberrations, making them indistinguishable from invasive tumors. The estimated time between STIC development and HGSOC progression is seven years ^14^, possibly leading to further molecular adaptations. Another explanation for the high proteomic similarity between STICs and ICs is that STICs do not represent precursor lesions but advanced tumors that retrogradely metastasize to the fallopian tube. However, given the lower frequency of this scenario ^14,15^, this hypothesis doesn’t explain the strong coherence of our proteomic data for the 32 matching STIC-invasive tumor pairs.

To further explore the proteomic heterogeneity of STICs, we integrated previously identified molecular subtypes from bulk transcriptomics and proteomics data ^2,7^, assigning nearly half of the epithelial samples to one predominant subtype. The inability to unambiguously assign all samples to one subtype might reflect HGSOC’s polyclonal nature and the co-existence of diverse tumor subpopulations with distinct gene expression programs in different spatial niches ^4,6,38^. Among the clearly assigned samples, we classified half as the proliferative subtype and half as immune-enriched, with a likely underlying differentiated tumor cell phenotype ^4^. While the clinical significance of HGSOC molecular subtypes remains debatable, our global proteomics data suggest a simple binary classification into two distinct epithelial proteotypes with therapeutic implications. Cluster 1 tumors showed an inflamed, immune-enriched signature with elevated interferon and TNF signaling, aligning with a recent spatial omics report ^62^ and high lymphocyte infiltration. Our findings generally agree with Kader et al.’s conclusion that interferon signaling from epithelial cells is an early occurrence in HGSOC development, but indicate this inflammatory characteristic is present in approximately half of all STICs. Proliferative cluster 2 samples showed low interferon-related signaling, were immune-deserted, and linked to higher patient age and likely more unfavorable clinical outcomes. This dichotomy is supported by long-known phenotypic differences in tumor-infiltrating CD3+ T cells, a favorable prognostic factor for roughly half of HGSOC patients ^63^. Our precise sampling through laser microdissection enabled us to disentangle distant from intra-epithelial immune cells, where only the latter marks tumors likely to respond to immune checkpoint inhibitors (ICI) ^20^. Notably, a comparable IFN-high immunophenotype (similar to cluster 1 tumors) associated with an up-regulation of antigen presentation and intra-epithelial immune cells was recently identified as a characteristic of metastatic deficient mismatch repair colorectal cancers responding to ICI treatment ^29^. Therefore, our deep proteomics data serve as a valuable resource to build new classifiers for improved patient stratification beyond PD-L1 assessment. This is clinically relevant since ICI treatment in ovarian cancer has yielded disappointing results ^20^, likely due to the absence of clear guidelines to tailor treatment to patients with the highest likelihood of therapeutic benefit.

The absence of an epithelial STIC proteotype distinct from concurrent invasive tumors contrasts with our stromal findings, which revealed a disease gradient of stromal remodeling from normal fallopian tube over STIC to invasive stroma. Stromal remodeling of STIC precursors featured both normal and invasive phenotypes, possibly reflecting different stages of progressive tumorigenic ECM remodeling. This suggests some STICs, despite high epithelial proteome similarity to invasive tumors, have not undergone complete stromal transformation. As all STICs in our cohort were accompanied by adjacent invasive tumors, it raises questions about the timescales of TME remodeling, clonal escape, and tissue invasion. While the presence or absence of stromal immune cell infiltration (e.g., CD8+ T cells) was strongly associated with the epithelial tumor proteotype (i.e., immune-enriched/differentiated [cluster 1] *versus* proliferative [cluster 2]), oncogenic ECM remodeling was a more uniform feature in our cohort. We found that one-third of the stromal proteome undergoes quantitative remodeling towards an immunosuppressive TME likely driven by myofibroblasts and M2-like macrophages. These changes include many known ECM degraders (cathepsins and MMPs), pro-inflammatory cytokines (TGFϕ31, TGFϕ32, IL16, and IL18), and many structural core matrisome proteins. On this basis, we extracted a 100-protein stromal signature of commonly upregulated and downregulated proteins. This signature features known disease drivers and oncogenic ECM proteins associated with ovarian cancer progression and metastasis (e.g., NNMT, CD163, and FN1) ^22,27,64^ and sheds light on proteins with high therapeutic potential. For example, thymidine phosphorylase (TYMP), a therapeutic target in metastatic colorectal cancer ^65^, showed a strong and gradual increase in the invasive stroma. Similarly, SYK, a non-receptor tyrosine kinase implicated in immune cell regulation ^66^, is targeted by Fostamatinib, which was approved by the FDA for the treatment of chronic immune thrombocytopenia ^67^. Notably, SYK also represents a promising strategy to target TAMs in pancreatic cancer ^68^, cells we found to be strongly enriched in the invasive stroma. Together, these data underscore the central role of the stromal compartment in HGSOC pathogenesis, the importance of tumor-stromal co-evolution, and the potential of the TME as a therapeutic target.

In summary, our study highlights the power of spatially and cell-type resolved proteomics to dissect the molecular underpinnings of early carcinogenesis and provides a rich proteomic resource for biomarker and drug target discovery.

## Limitations

One limitation of our study is that we cannot be certain all analyzed STICs are true precursor lesions or represent metastasis to the fallopian tube epithelium. However, the latter scenario is less frequently observed according to two recent genomics studies ^14,15^. Currently, no established histopathological criteria differentiate between these scenarios ^69^. To avoid collecting retrograde metastases, we sampled only intraepithelial, non-invasive TP53 mutant neoplastic epithelium, not free-floating cells or detached cell clusters in the tubal lumen ^70^. To exclude retrograde colonization entirely, collection of STICs from patients who have undergone prophylactic risk-reducing salpingo-oophorectomy is necessary (fallopian tube and ovary resection). However, this alternative would entail constraints. The number of samples would be limited as the prevalence of STICs in risk-reducing surgery ranges between 0 and 25% ^71^. The samples would also lack concurrent invasive tumors, limiting analysis to normal fallopian tube and STIC regions. Our patient cohort would exclude the majority of sporadic HGSOC, instead including younger individuals with germline *BRCA1/2* mutations.

In addition to STICs, other fallopian tube lesions linked to HGSOC are secretory cell outgrowths (SCOUTs), p53 signature lesions, and serous tubal intraepithelial lesions (STILs). These are considered precursors of STIC, and ultimately HGSOC. Except for two lesions reclassified as STILs, this study did not examine the full range of lesions representing the transition between normal epithelium and STIC. Their clinical relevance is unclear, and future studies are needed to dissect the full spectrum of HGSOC development through various precursor lesions to characterize the proteomes of SCOUTs, p53 signature lesions, and STILs. Nevertheless, we believe the applied methodology and acquired deep spatial proteomic data provide an excellent starting point and reference dataset to understand the earliest proteomic changes and underlying biology of this devastating disease.

## Acknowledgments

We thank our colleagues at the Max Delbrück Center (MDC) and Charité for their support and fruitful discussion. In particular, Janett König supported multiplex imaging experiments. We also thank Tobias Janik and Ines Koch for their help with slide preparation and cell line experiments. Furthermore, we acknowledge the MDC technology platform ‘Proteomics’ and ‘Advanced light microscopy’ for their great support. Gaetano Gargiulo (MDC) and Elena Ioana Braicu (Charité) we thank for their critical feedback on the manuscript and Gina Dörpholz and Pia Larsen for administrative support.

A.M, S.F., and F.C. acknowledge funding support by the Federal Ministry of Education and Research (BMBF), as part of the National Research Initiatives for Mass Spectrometry in Systems Medicine, under grant agreement No. 161L0222. This project received funding from the European Research Council (ERC) under the European Union’s Horizon 2020 research and innovation program (grant agreement No. 101115681) and support by the ERC (ERC starting grant). M.P.D. is a Clinician Scientist as part of the Berlin Institute of Health Clinician Scientist Program. The work of M.P.D. is supported by a DKTK Berlin Young Investigator Grant 2022, and Berliner Krebsgesellschaft Grant (DRFF202204).

## Author contributions

Conceptualization, A. M., M.P.D., E.T.T., F.C.; methodology, A.M., M.P.D., S.F., F.C.; experiments, A.M., M.P.D.; data curation, A.M., M.M., W.D.S; data analysis, A.M., M.P.D., and F.C.; figures, A.M., M.P.D., and F.C.; supervision, E.T.T., F.C.; funding acquisition, M.P.D., E.T.T., F.C.; writing the original draft, F.C. All authors reviewed and edited the manuscript.

## Methods

### RESOURCE AVAILABILITY

#### Lead contact

Further information and requests for resources and reagents should be directed to and will be fulfilled by the lead contact, Fabian Coscia.

#### Materials availability

This study did not generate new materials.

#### Data and code availability

The mass spectrometry proteomics data have been deposited to the ProteomeXchange Consortium (http://proteomecentral.proteomexchange.org) via the PRIDE partner ^72^ with the dataset identifier PXD060547. All data reported in this paper will be shared by the lead contact upon reasonable request. Any additional information required to reanalyze the data reported in this paper is available from the lead contact upon request.

## EXPERIMENTAL MODEL AND SUBJECT DETAILS

### Sample collection and patient cohort

We retrieved all samples containing the term “STIC” and/or “serous tubal intraepithelial carcinoma” in the pathology report from the pathology archive of the Institute of Pathology, Charité, between 2013 and 2022. Two pathologists (M.P.D. and E.T.T.) performed two independent rounds of reclassification of the precursor lesions according to the criteria proposed by Vang *et al*.^23^, using three whole slides stained with H&E, p53, and Ki67. Briefly, STIC was defined by a lesion morphologically suspicious or unequivocal for STIC and an immunohistochemical p53 aberrant expression and a proliferation rate (Ki67) > 10%. Only samples in which full agreement was reached were included in the proteomic analysis. In addition, we excluded all samples that received chemotherapy prior to the resection of tubal lesions. Clinical data were obtained from the Tumour Bank Ovarian Cancer Network (www.toc-network.de) or the Charité Comprehensive Cancer Center (https://cccc.charite.de). This study was approved by the local ethics committee (EA1/110/22).

#### Cell line models

The human ovarian carcinoma cell line OAW-42 was obtained from the European Collection of Animal Cell Cultures (Salisbury, United Kingdom). OVCAR-8 was obtained from the laboratory of Ernst Lengyel (Department of Obstetrics and Gynecology/Section of Gynecologic Oncology, University of Chicago, Chicago, IL, USA). ES-2 was obtained from the American Type Culture Collection and EFO-21 was obtained from Dr. Fritz Hölzel (Department of Gynecology, University Hospital Eppendorf, Hamburg, Germany). The cells were cultured in DMEM (Gibco, #21885-025), all supplemented with 10% fetal bovine serum (Capricorn, #FBS-16A), no added antibiotics, at 37°C with 5% CO2 and 95% humidity. Prior to the study, the cytogenetic analysis and cell authentication of the cells was performed at the DNA-Fingerprinting Facility at Charité Berlin using short tandem repeat DNA. All cell lines were tested for mycoplasma contamination using PCR mycoplasma kit (Biontex, #M030/050).

## METHOD DETAILS

### Cell viability assay

Cell viability assay was performed in 96 well-plates. For OAW-42, OVCAR-8, and ES-2 we plated 4000 cells/well, and for EFO-21 we plated 6000 cells/well and we cultured the cells in full growth medium for 24 h. The medium was then removed and replaced with new full medium containing different concentrations of AY-9944 (MedChemExpress, #HY-107420), carboplatin (Merck, #C2538) or combinations of the two drugs. The plates were incubated at 37 °C, 5% CO_2_ for 72 h. After the treatment, 11 µL of MTT assay reagent (Merck, #M2128) was added to the medium, and incubated at 37 °C, 5% CO_2_ for 4 h. Following MTT incubation, the media and MTT were removed, 100 μL of DMSO was added to all wells, including controls and spectrophotometric absorbance of the samples was detected by using a microplate spectrophotometer (BioTek, Synergy 2) at 540 nm wavelength.

#### Homologous Repair Deficiency Analysis

HRD analysis was performed using the (Northeastern German Society for Gynecologic Oncology) NOGGO GIS v1 Assay, as previously described ^73^. Briefly, from tumor rich regions of invasive carcinoma from ten to twenty sequential 5 micrometer thick FFPE slides DNA was extracted. 50 to 100 ng of tumor DNA was used for library preparation using hybrid capture XT HS2 chemistry (Agilent Technologies) targeting all exonic bases as well as a minimum of 10 bp flanking region of 57 genes, including 35 HRR genes, as well as selected driver genes and more than 20,000 genome-wide evenly distributed single nucleotide polymorphism (SNP) loci that enable the detection of allele-specific copy number alterations (CNA). Next, the libraries were subjected to sequencing with a minimum of 20 million reads using 100 bp read length on a NextSeq 2000 instrument (Illumina, San Diego, CA, USA). The genomic instability score (GIS) was calculated using an allele-specific copy number profile and three measures of HRD based on the PureCN output: percent loss of heterozygosity (PLOH), percent copy number alteration (PCNA), and percent telomeric copy number alteration (PTCNA). A GIS cutoff of “83” or the presence of a pathogenic mutation in either BRCA1 or BRCA2 was used to define HRD-positive cases. Mutation calling was conducted using SEQUENCE Pilot Software, Version 5.4.0 (JSI Medical Systems GmbH, Ettenheim, Germany).

#### Immunohistochemistry (IHC)

Immunohistochemical staining was performed on a BenchMark XT immunostainer (Ventana Medical Systems, Tucson, AZ, USA). For antigen retrieval, sections were incubated in CC1 mild/standard buffer (Ventana Medical Systems, Tucson, AZ, USA) for 30 min at 100 °C. The sections were stained with anti-Ki67 antibody (M7240, Dako, 1:50, CC1 mild buffer), anti-p53 (M7001, Dako, 1:50, CC1 standard buffer), anti-PAX8 (760-4618, Roche/Ventana, ready to use, CC1 mild buffer), and anti-p16 (805-4713, Roche/Ventana, 1:2, CC1 mild buffer) for 60 min at room temperature, and visualized using the avidin–biotin complex method and DAB. We stained the cell nuclei by additionally incubating for 12 min with hematoxylin and bluing reagent (Ventana Medical Systems, Tucson, AZ, USA). Histological images were acquired with a PANNORAMIC 1000 digital slide scanner (3DHISTECH).

| Antibody | Company | Catalog # | Clone | Species | Dilution | Pre-preparation |
| --- | --- | --- | --- | --- | --- | --- |
| p53 | Dako/Agilent | M7001 | DO-7 | Mouse | 1:50 | CC1 standard |
| PAX8 | Roche/Ventana | 760-4618 | MRQ-50 | Mouse | ready to use | CC1 mild |
| Ki67 | Dako/Agilent | M7240 | MIB-1 | Mouse | 1:50 | CC1 mild |
| p16 | Roche/Ventana | 805-4713 | E6H4 | Mouse | 1:2 | CC1 mild |

#### Cyclic immunofluorescence (CyCIF) staining and imaging

Prior to immunostaining, tissue sections were incubated at 60 °C for 30 min, followed by deparaffinization and sequential rehydration as follows: two 5-minute immersions in neo-clear buffer, two 2-minute washes in 99% ethanol, and sequential 2-minute washes in 80%, 70% ethanol, and 1x PBS (twice for 1 min each). Heat-mediated antigen retrieval was performed in Tris-EDTA (pH 9) using a steamer for 25 min, followed by cooling to room temperature in the retrieval solution. Slides were then washed three times in 1x PBS.

To reduce tissue autofluorescence, sections were pre-bleached for 30 min under direct white light in 4.5% H_2_O_2_ and 24 mM NaOH diluted in 1x PBS. Following three additional PBS washes, tissue sections were outlined with a PAP pen to minimize the reaction volume and blocked with 3% BSA in 1x PBS for 30 min at room temperature. Antibody incubation was performed overnight at 4 °C in a blocking buffer in a humidified staining chamber. Immunofluorescence staining was conducted over several cycles. Except for p53, all antibodies were directly conjugated (listed below). For p53 staining, sections were washed in 1x PBS after primary antibody incubation and then incubated with a secondary fluorescently conjugated antibody (A555 donkey anti-mouse, 1:500) at room temperature for 1 h. Slides were subsequently washed in 1x PBS, counterstained with Hoechst 33342 (1 µg/mL) for 5 min at room temperature, washed again, mounted with 10% glycerol in 1x PBS and imaged. Following imaging, coverslips were removed by soaking the slides in 1x PBS within a vertical Coplin jar on a platform shaker. Detached slides were washed in 1x PBS, bleached for 30 min under direct white light in 4.5% H_2_O_2_ and 24 mM NaOH diluted in 1x PBS, and washed again in 1x PBS before proceeding to the next round of primary antibody staining. This process was repeated twice to achieve staining for 10 markers. After the final cycle, coverslips were removed by soaking the slides in 1x PBS within a vertical Coplin jar on a platform shaker. Once detached, the slides were rinsed with Milli-Q water, air-dried, and stored at 4 °C. Imaging was conducted using a Zeiss Axioscan 7 slide scanner equipped with the Colibri 7 LED light source and an EC Plan-Neofluar 20x/0.50 M27 objective at 2×2 binning. Stitching of the raw images was performed using ZEN software (version 3.5, Blue Edition), with the DAPI channel set as the reference for all channels and the following parameters: minimal overlap of 5%, maximal shift of 15%, Comparer set to Optimized, and Global Optimizer set to *Best*.

| Target | Dilution | Fluorophore | Source | Order no. | RRID |
| --- | --- | --- | --- | --- | --- |
| Ki67 | 1:100 | Alexa Fluor 488 | Cell Signaling | 11882 | AB_2687824 |
| p53 | 1:25 | - | Agilent Dako | M7001 | AB_2206626 |
| anti-rabbit | 1:500 | Alexa Fluor 555 | Invitrogen | A32794 | AB_2762834 |
| CD4 | 1:50 | Alexa Fluor 647 | Cell Signaling | 40568S | AB_3492108 |
| CD8 | 1:100 | Alexa Fluor 488 | Cell Signaling | 79594S | AB_2799935 |
| FOXP3 | 1:50 | PE | Biolegend | 320107 | AB_492986 |
| CD11c | 1:50 | Alexa Fluor 750 | BioTechne | NBP2-54432AF750 | AB_3083691 |
| CD163 | 1:50 | Alexa Fluor 488 | Abcam | ab218293 | AB_2889155 |
| panCK | 1:100 | eFluor™ 570 | Thermofisher | 41-9003-82 | AB_11218704 |
| PAX8 | 1:50 | CoraLite 647 | Proteintech | CL647-10336 | AB_2920213 |
| COL1A1 | 1:50 | Alexa Fluor 750 | BioTechne | NB600-408AF750 | AB_10000511 |

#### Image analysis and contour export for laser microdissection

QuPath (version 0.4.3) was utilized for conducting image analysis. Regions of interest were manually annotated in QuPath following the image analysis process. To ensure accurate contour transfer between the screening and laser microdissection microscopes, three tissue reference points (x-y coordinates) were selected. The contours and reference points were then exported in geojson format and converted into .XML format, which is compatible with Leica LMD7 software. The shape processing code can be accessed at github.com/CosciaLab/Qupath_to_LMD, and it employs geopandas (Version 0.12.2) and py-lmd ^24^ (Version 1.0.0).

#### Laser microdissection

For laser microdissection, three slides were prepared for each sample and mounted on PPS-Membrane Frame Slides (Leica). The slides were then stained immunohistochemically with p53 (5 µm thick slide), PAX8 (2.5 µm thick slide), and Ki67 (2.5 µm thick slide). Whenever feasible, the specimen was collected from the p53 stained slide, while the other slides served mainly for orientation. Sample annotation was conducted under the direct supervision of at least one of the study pathologists (M.P.D./E.T.T.).

For laser microdissection-based tissue collection, we employed the Leica LMD 7 system along with Leica Laser Microdissection V 8.3.0.08259 software. The tissue was cut using a 20x objective in either brightfield or fluorescence mode. We applied the following laser parameters for the 20x objective (HC PL FL L 20x/0.40 CORR): 56 power, 1 aperture, 15 speed, 1 middle pulse count, -1 final pulse, 37 – 45% head current (adjusted based on tissue type and section thickness), 801 pulse frequency, and 101 offset. The cut contours were then collected into a low-binding 384-well plate (Eppendorf 0030129547), which was set up using the ’universal holder’ function with a single empty well separating each sample.

#### Sample preparation for MS-based proteomic analysis

Automated cutting was employed to gather tissue specimens following contour import into 384-well plates with low binding properties. To ensure tissue settled at the well bottoms post-LMD collection, each well received 15 µL of acetonitrile, underwent brief vortexing, and was vacuum dried (15 min at 60 °C). A subsequent well inspection was conducted prior to proteomics sample preparation to verify successful collection. The DDM-based protocol utilized a lysis buffer comprising 0.025% DDM, 5mM TCEP, 20mM CAA, and 0.1M TEAB in water. A MANTIS Liquid Dispenser with high-volume diaphragm chips was used to add 4 μl of lysis buffer to each sample well. The plate was sealed with PCR ComfortLid and heated at 95 °C for 60 min. After brief cooling, 1 µL of LysC (10 ng/ µL in 0.1M TEAB [pH 8.5] and 30% ACN in milliQ water) was introduced, followed by digestion for at least 2 h at 37 °C in a thermal cycler (50 °C lid temperature). Next, 1 µL of trypsin (10 ng/μl containing 10% ACN and 0.1M TEAB [pH 8.5] in milliQ water) was added, with overnight incubation at 37 °C in the thermal cycler. The following day, digestion was stopped by the addition of trifluoroacetic acid (TFA, final concentration 1% v/v), and samples underwent vacuum drying before peptide clean-up.

#### Peptide clean-up with C18 tips

Peptide purification was conducted using Evotips (Evosep, Odense, Denmark) following the manufacturer’s instructions. The tips were prepared by adding 20 μl of buffer B (99.9% ACN, 0.1% FA) to each C-18 tip (EV2013, Evotip Pure, Evosep) followed by centrifugation at 700 rpm for 1 min. Subsequently, 20 μl of buffer A (99.9% water, 0.1% FA) was added to the top of each C-18 tip, which was then activated in isopropanol for 20 s and centrifuged again at 700 rpm for 1 min. Peptides were then applied to the Evotips, washed once with 20 μL of buffer A, and finally loaded with 200 μL of buffer A. The tips were also submerged at the bottom in buffer A before initiating the LC-MS analysis.

#### MS-based proteomic analysis

Liquid chromatography was performed using the Evosep One LC system (Evosep, Odense, Denmark) connected to a trapped ion mobility spectrometer with quadrupole time-of-flight mass spectrometer (timsTOF Ultra, Bruker Daltonik, Bremen, Germany) with a nano-electrospray ion source (CaptiveSpray, Bruker Daltonik, Bremen, Germany). Digested peptides were loaded on the Evosep Performance column (EV1137, 150 µm inner diameter, packed with 1.5 µm C18 beads) at 40 °C. Chromatographic separation was performed using an Evosep 30SPD gradient. The solvents utilized were LC-MS-grade water containing 0.1% formic acid (buffer A) and acetonitrile with 0.1% formic acid (buffer B). For the dia-PASEF analysis, we employed a method comprising eight dia-PASEF scans divided into three ion mobility windows per scan. This covered a mass-to-charge ratio range of 400-1000 m/z using 25 Th windows and an ion mobility range from 0.64 to 1.37 Vs cm^-2^. The mass spectrometer was run in high sensitivity mode, with accumulation and ramp time set to 100 ms, and capillary voltage at 1750V. The collision energy was configured as a linear ramp, starting at 20 eV at 1/k_0_ = 0.6 Vs cm^-2^ and increasing to 59 eV at 1/k_0_ = 1.6 Vs cm^-2^. This collision energy ramp was applied linearly as a function of ion mobility, decreasing from 59 eV at 1/k_0_ = 1.6 Vs cm^-2^ to 20 eV at 1/k_0_ = 0.6 Vs cm^-2^.

#### Mass spectrometry raw file analysis

For the analysis of dia-PASEF raw files and generation of spectral libraries, we employed DIA-NN (version 1.8.1). The human FASTA file was obtained from the UniProt database (2022 release, UP000005640_9606, downloaded on April 8th 2022). To generate in silico predicted libraries, we provided the human FASTA file along with commonly encountered contaminants^74^ . We enabled deep learning-based predictions for spectra, RTs, and IMs within the 300-1200 m/z mass range. Fixed modifications included N-terminal M excision and cysteine carbamidomethylation. We allowed up to two missed cleavages and set the precursor charge to 2–4. DIA-NN was run in default mode with slight modifications. We configured MS1 and MS2 accuracies to 15.0, set scan windows to 0 (for DIA-NN assignment), enabled isotopologues, activated match-between-runs, applied heuristic protein inference, and disallowed shared spectra. Protein inference was based on genes. We set the neural network classifier to single-pass mode and selected ’Robust LC (high precision)’ as the quantification strategy. Cross-run normalization was configured as ’RT-dependent,’ library generation as ’smart profiling,’ and speed and RAM usage as ’optimal results.’

## QUANTIFICATION AND STATISTICAL ANALYSIS

### Proteomic data analysis

Proteomic data analysis was performed within the R environment (https://r-project.org/version 4.3.2) with the following packages: tidyverse (version 2.0.0), Rstatix (version 0.7.2), UpSetR package (version 1.4.0), FactoMineR (version 2.11), factoextra (version 1.0.7), ggpubr (version 0.6.0), corrplot (version 0.92), ComplexHeatmap (version 2.18.0), RColorBrewer (version 1.1.3), circlize (version 0.4.16), dendsort (version 0.3.4), ggcorrplot (version 0.1.4.1), ggplot2 (version 3.5.1), ggrepel (version 0.9.5), viridis (0.6.5) and networkD3 (version 0.4). Perseus ^75^ (version 1.6.15.0) was used for additional exploratory data analysis. Prior to statistical testing, data were first filtered to keep only proteins with 70% non-missing values in at least one group, or more stringently with 70% in all groups. Missing values were imputed based on a normal distribution (width = 0.3, downshift=1.8). Pairwise comparisons were computed using a two-sided Student’s t-test. For multiple group comparisons, analysis of variance (ANOVA) was used. For both tests, a permutation-based FDR of 5% was applied to correct the multiple hypothesis testing. Pathway enrichment analysis was performed in Perseus based on Fisher’s exact test (categorical data) or 1D pathway enrichment analysis ^76^. Hallmark gene sets, WikiPathways, and the Reactome pathways were enriched terms filtered using a Benjamini-Hochberg FDR cut-off of 0.05. The minimum category size was set to five. The annotation matrix algorithm of Perseus (version 1.5.0.3) was additionally employed to globally compare pathway-level changes across multiple samples (**Supplementary** Fig. 3B). This algorithm tests the difference in any protein annotation from the overall intensity distribution of the sample. The resulting pathway scores were z-scored for relative comparison.

For subtype classification of high-grade serous ovarian cancers based on TCGA subtypes, the consensusOV package (version 1.26.0, get.consensus.subtypes function) was used. The package calculates probability for each subtype and assigns each sample to the subtype with the highest probability. To characterize the previously identified fallopian tube epithelial cell types in the dataset, we used the signature matrix from Dinh *et al*. ^34^. For the systematic analysis of ECM changes, the matrisome database described by Renner *et al*. ^44^ was used.

To prioritize proteins upregulated in the invasive stroma during HGSOC progression (Fig. 5), we employed support vector machine (SVM) classification using the Perseus software. Before classification, data were first filtered (70% in at least one group), log2-transformed, and imputed based on a normal distribution (width = 0.3, downshift = 1.8). The SVM was then trained on the stromal proteomes to determine the optimal classification parameters. To achieve the lowest classification error, we systematically tested kernels and the corresponding parameters. We used Radial basis function (RBF) kernel, parameter sigma (σ) = 5 and parameter C =10. For cross validation, the “leave-one-out” function was used, and ANOVA was used as the feature ranking method with an s*_0_* value ^77^ of 0.1. The optimal number of discriminating protein features was then determined from the error percentage curve, which resulted in an error rate of 5% for the top 100 features.

The list of FDA-approved drug targets was retrieved from the human protein atlas database (https://www.proteinatlas.org/search/protein_class:FDA+approved+drug+targets).

For drug-response curve and drug synergy calculations, we used the SynergyFinder R package and web application ^78^. Three reference models were used to assess drug synergy based on the highest single agent (HSA) score, Bliss independence (Bliss) and zero interaction potency (ZIP) ^79^.

### Declaration of generative AI and AI-assisted technologies in the writing process

During the preparation of this manuscript, the authors used Paperpal in order to improve readability and language of the text. After using this tool, the authors reviewed and edited the content as needed and take full responsibility for the content of the publication.

---

**Figure S1.**
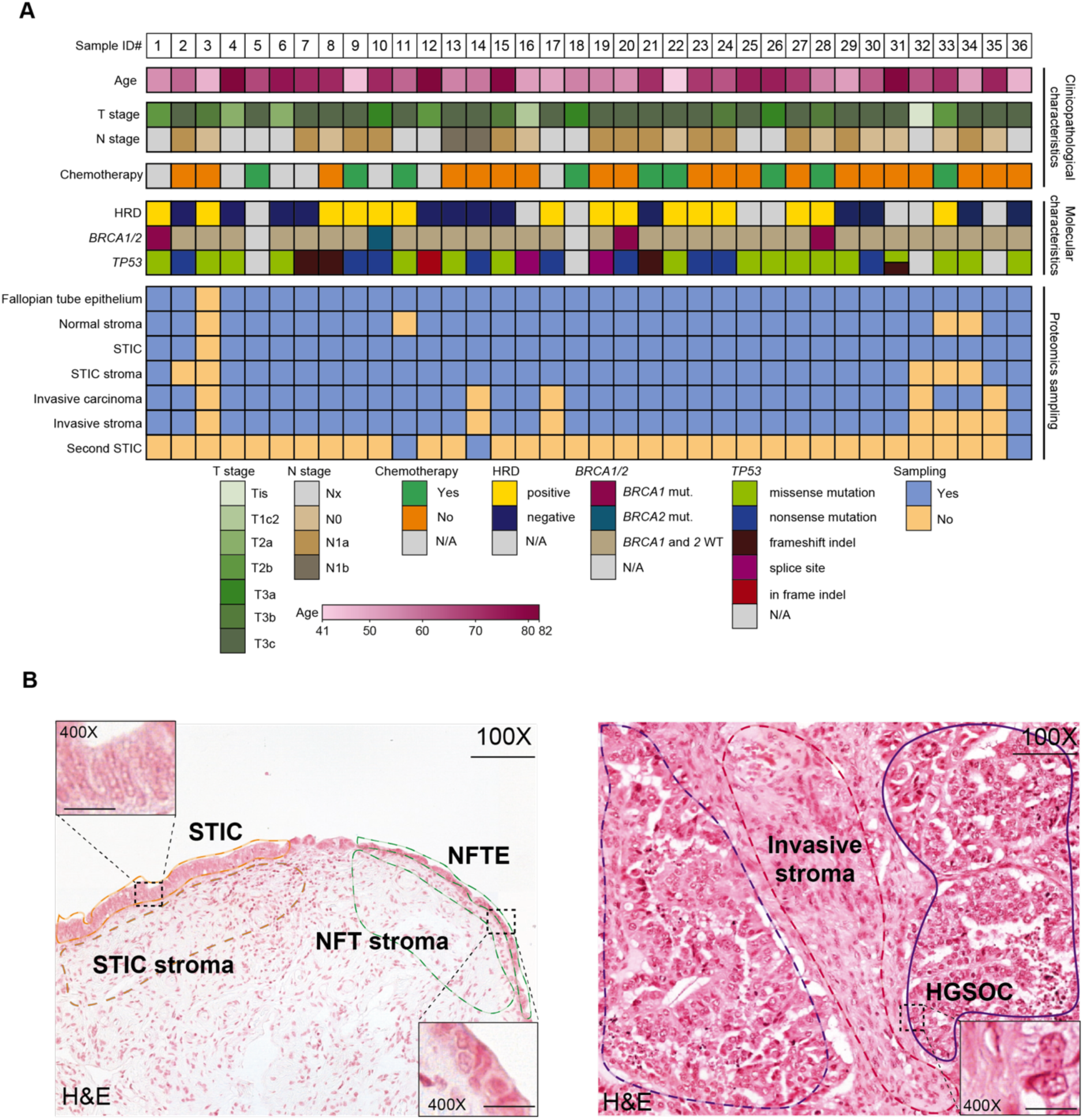
Pathology-guided ultra-low input proteomics of HGSOC precursor lesions. **(A)** Patient cohort (*n* = 36) with clinicopathological and molecular characteristics and proteomic sampling. **(B)** Representative H&E stains of normal fallopian tube epithelium (NFTE), serous tubal intraepithelial carcinoma (STIC), and invasive carcinoma (IC) and corresponding epithelial and stromal compartments. Scale bars: 100X: 100 *μm*, 400X: 20 *μm*.

**Fig. S2:**
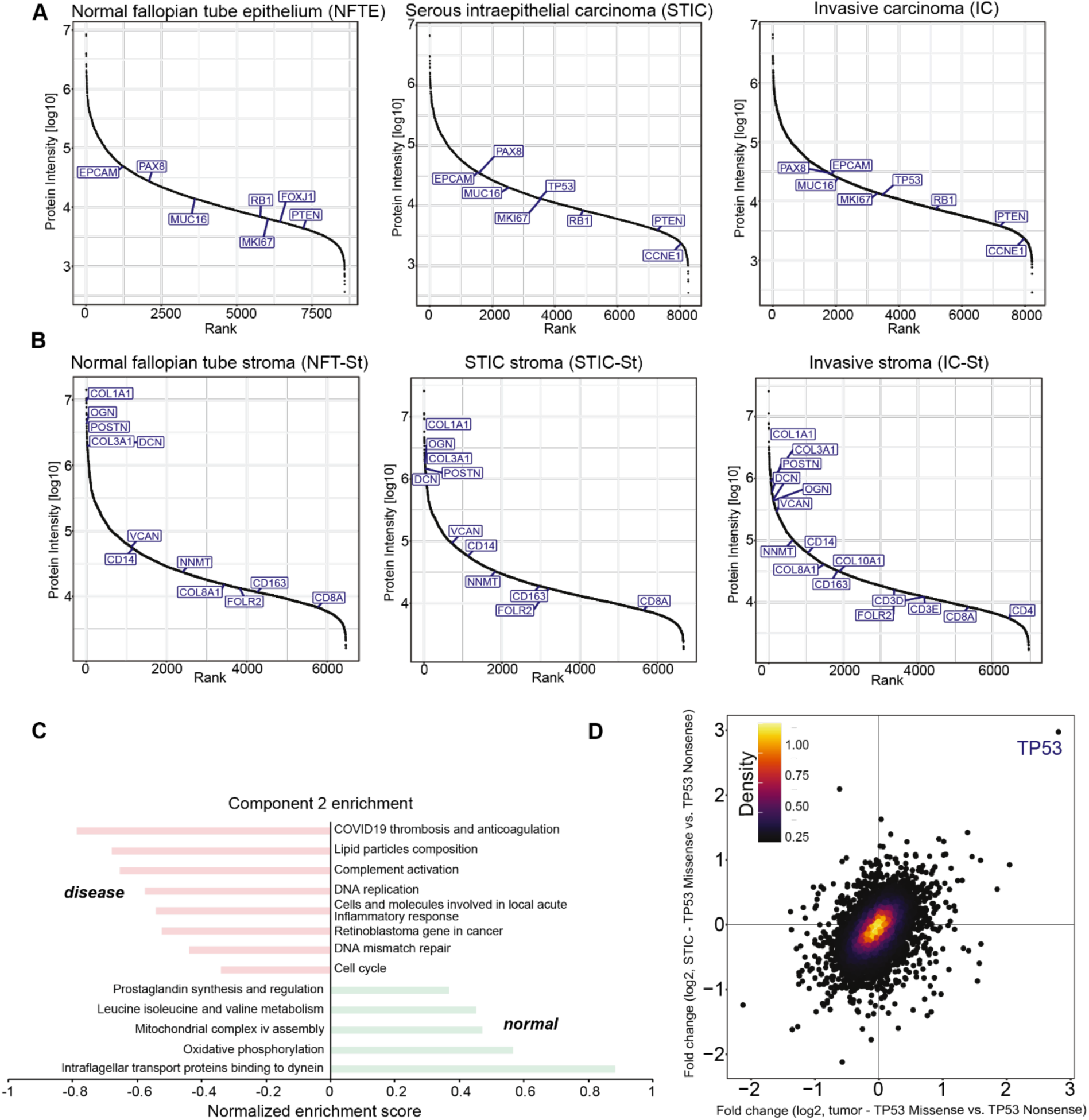
Deep spatial proteomics quantifies disease-specific alterations at bulk level resolution. **(A and B)** Dynamic range of median protein abundance for epithelial and stromal compartment samples, respectively. Known ovarian cancer-, cell type-, and stromal markers are highlighted. A minimum of 18 quantified values were required for each marker to be displayed. Proteins with 50% valid values for each group are shown. **(C)** Pathway enrichment analysis (Hallmarks) based on PC2 revealing differences between normal and disease compartments. Selected terms with a Benjamini-Hochberg FDR < 0.05 are shown. **(D)** Scatter plot of t-test results between p53 overexpression and p53-null mutation patients, indicating that p53 is the only differentially abundant protein between the two groups.

**Fig. S3.**
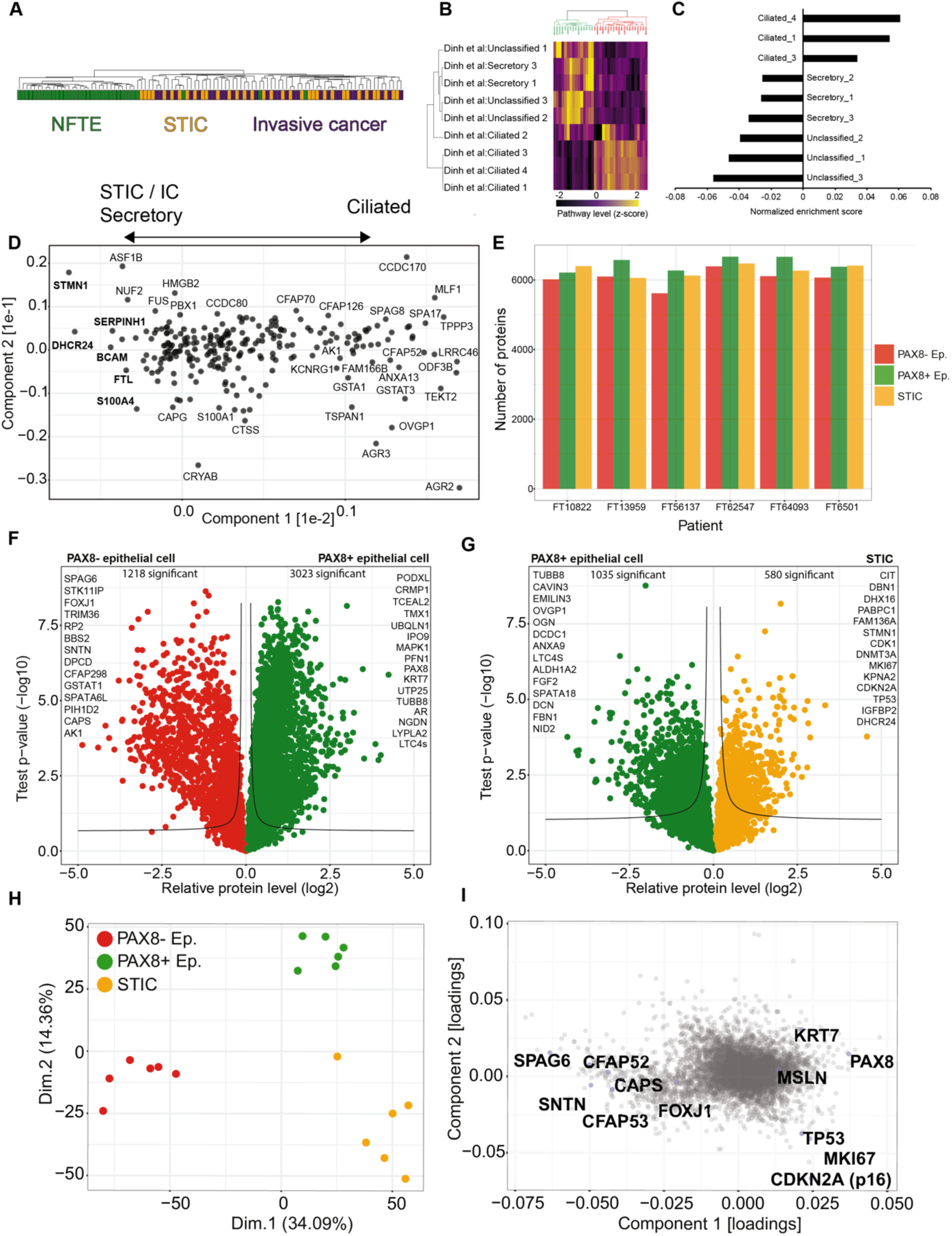
STICs and invasive tumors feature strong cell-of-origin signatures. **(A)** Hierarchical tree showing the clustering pattern across regions (normal fallopian tube epithelium, STIC, and invasive fallopian tube carcinoma), revealing three epithelial samples that are pre-malignant lesions based on proteome profiling. **(B)** Unsupervised hierarchical clustering of all normal fallopian tube epithelial samples based on cell type abundance scores. Cell type abundance scores were calculated based on the annotation matrix algorithm (Perseus) for all fallopian tube epithelial cell types identified by Dinh *et al*. Two main clusters of secretory and ciliated cell-enriched normal fallopian tube epithelium samples were identified. (**C**) Enriched cell type signatures along PC1, related to Fig. 3F. **(D)** PCA loadings related to Fig. 3F. Note that proteins on the left are higher in STIC, invasive carcinoma, and secretory-like epithelial cells, whereas proteins on the right are higher in ciliated epithelial cells**. (E)** Barplots showing the number of proteins identified in the PAX8 negative, PAX8 positive, and STIC compartments across six patients. **(F)** Volcano plot of the pairwise proteomic comparison between PAX8 positive (green) and PAX8 negative (red) epithelial samples. Cell and functional markers with the highest abundance change and significance are highlighted (two-sided Student’s t-test, FDR < 0.05). **(G)** Volcano plot of pairwise proteomic comparison between STIC (yellow) and PAX8 positive (green) samples. Cell and functional markers with the highest abundance change and significance are highlighted (two-sided Student’s t-test, FDR < 0.05). **(H)** PCA of all quantified proteins comparing STICs, secretory, and ciliated cell samples. **(I)** PCA loadings show known secretory and ciliated markers. Related to panel (H).

**Fig. S4.**
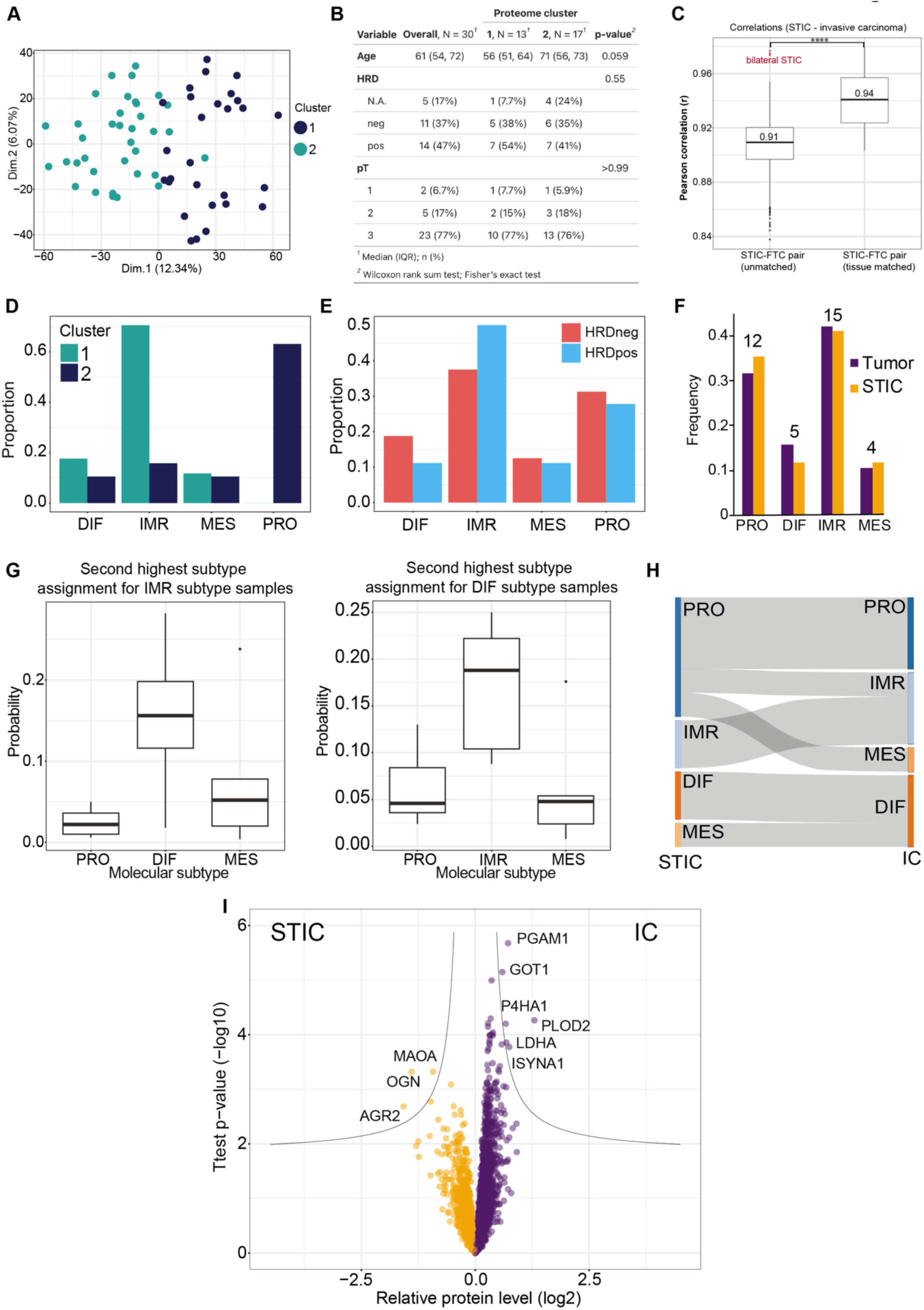
High proteome similarity between STICs and invasive fallopian tube cancer. **(A)** Principal component analysis of STIC and invasive carcinoma samples corresponding to figure 3A. **(B)** Multivariate statistical analysis of patients in clusters 1 and 2. Note that cluster 2 samples showed a trend towards higher patient age at diagnosis (*p* = 0.059). **(C)** Boxplots showing proteomic correlations between STIC and invasive carcinoma samples between matched and unmatched samples, corresponding to intrapatient and interpatient correlations. Boxplots define the range of the data (whiskers), 25th and 75th percentiles (box), and medians (solid line). **(D)** Molecular subtype assignment of cluster 1 and 2 samples based on the consensusOV algorithm^37^. Only samples with a margin score of >0.2 were included for high classification confidence. **(E)** Comparison between molecular subtypes and HRD status. **(F)** Frequency of molecular subtypes of STIC and IC proteomes. **(G)** Boxplots indicating the second highest molecular subtype classification probabilities for samples with immunoreactive and differentiated subtype calls. Boxplots define the range of the data (whiskers), 25th and 75th percentiles (box), and medians (solid line). **(H)** Sankey plot showing convergence or divergence of molecular subtypes for patient-matched STIC and invasive carcinoma pairs. **(I)** Volcano plot of pairwise proteomic comparison between STIC (yellow) and invasive fallopian tube carcinoma (violet) samples. Markers with the highest abundance change and significance are highlighted (two-sided Student’s t-test, FDR < 0.05).

**Fig. S5:**
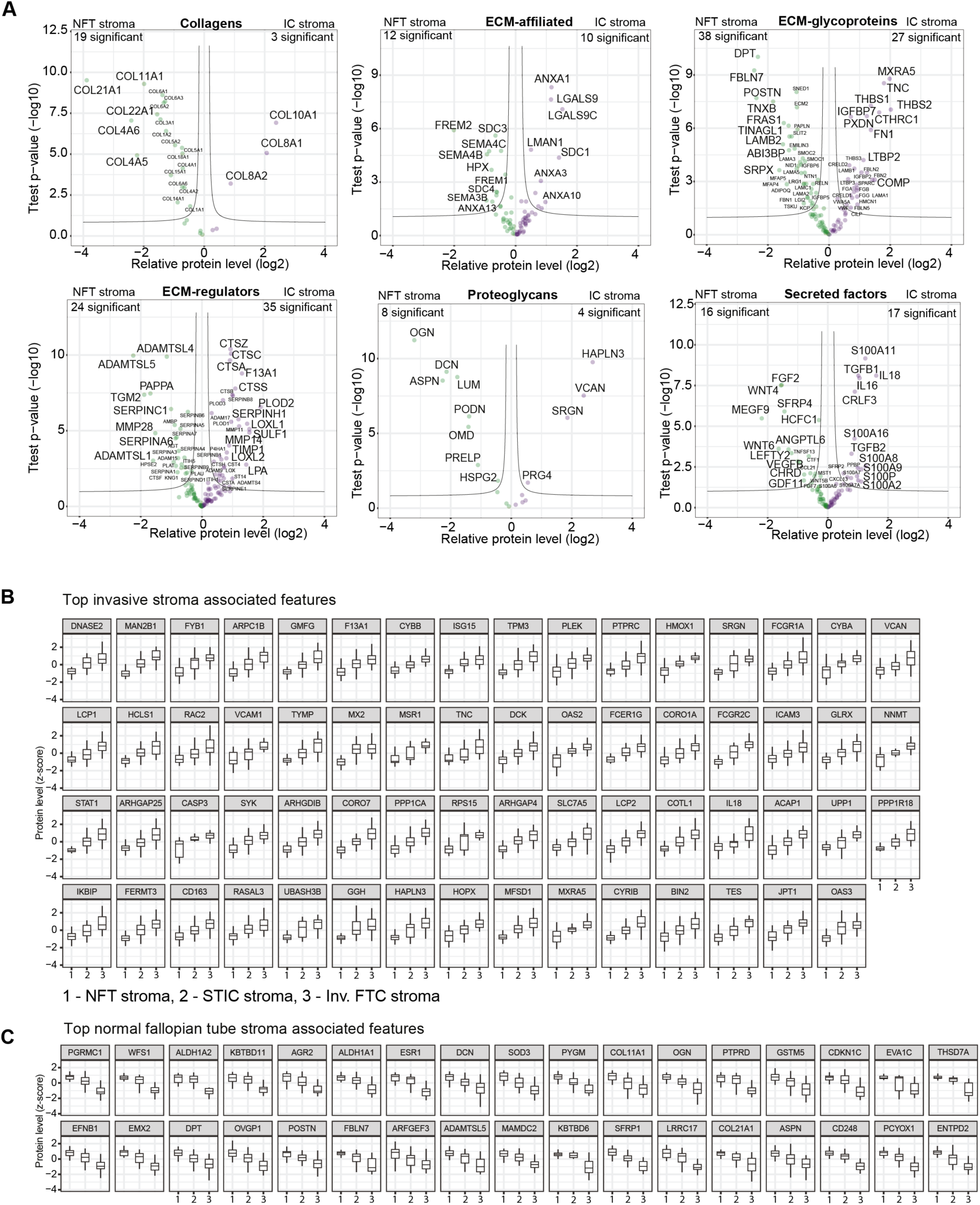
Progressive ECM remodeling and co-evolution of an immunosuppressive microenvironment. **(A)** Boxplots of individual relative protein levels (group average, z-score) for inv. FTC stroma-associated features. Boxplots define the range of the data (whiskers), 25th and 75th percentiles (box), and medians (solid line). **(B-C)** Boxplots of relative protein levels (group average, z-scored) for invasive stroma (B) and normal stroma (C)-associated features. Top-ranked protein features were identified by support vector machine classification, as shown in Fig. 5E. Boxplots define the range of the data (whiskers), 25th and 75th percentiles (box), and medians (solid line).

**Fig. S6:**
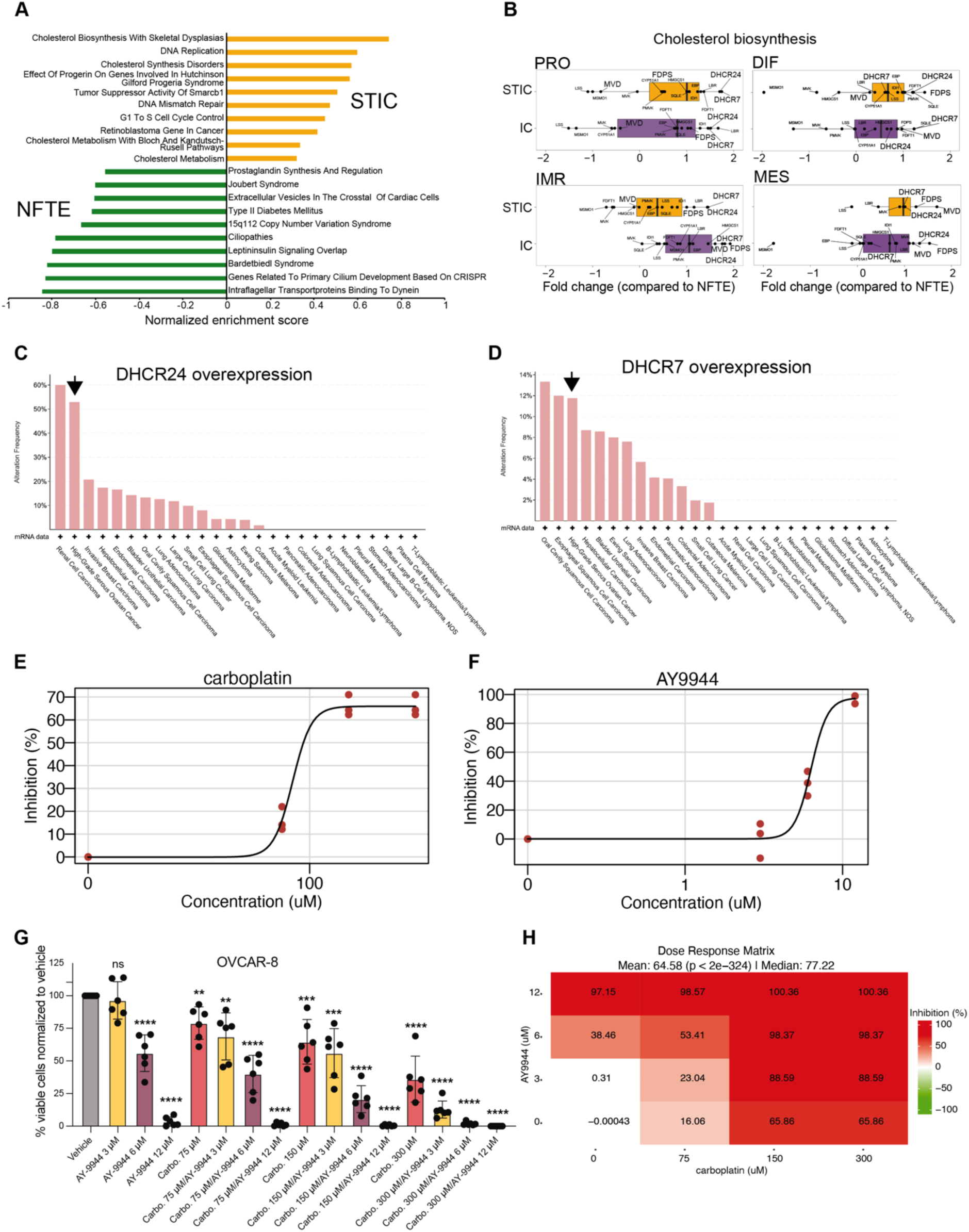
Identification of commonly dysregulated pathways of therapeutic relevance. **(A)** Pathway enrichment analysis (Hallmarks) based on t-test difference between STIC (yellow) and invasive fallopian tube carcinoma (violet) samples. Selected pathways with a Benjamin-Hochberg FDR < 0.05 are shown. **(B)** Protein fold changes (STIC vs. NFTE and IC vs NFTE, FDR < 0.05) of all cholesterol biosynthesis pathway proteins across 4 molecular subtypes of HGSOC. Proteins with significant fold change are highlighted in bold. **(C and D)** Frequency of DHCR24, DHCR7 genetic alteration (overexpression) respectively across different cancer types analyzed by cBioPortal. **(E)** Dose-response curve of OVCAR-8 cells treated with different carboplatin concentrations. (**F)** Dose-response curve of OVCAR-8 cells treated with different concentrations of the DHCR7 inhibitor AY9944 **(G)** Proliferation assay matrix of OVCAR-8 treated with different combinations of carboplatin and the DHCR-7 inhibitor (AY-9944). Experiments were performed in biological sextuplicate (with n = 3 technical replicates each). The same results are shown as bar plots. Mean ± SD; significance test: unpaired Student’s t-test: ns (not significant), ∗∗p < 0.01, ∗∗∗p < 0.001, ∗∗∗∗p < 0.0001. **(H)** Dose-response matrix of OVCAR-8 cells treated with different doses of carboplatin and AY-9944.

